# Feasibility of decoding covert speech in ECoG with a Transformer trained on overt speech

**DOI:** 10.1101/2024.02.05.578911

**Authors:** Shuji Komeiji, Takumi Mitsuhashi, Yasushi Iimura, Hiroharu Suzuki, Hidenori Sugano, Koichi Shinoda, Toshihisa Tanaka

## Abstract

Several attempts for speech brain–computer interfacing (BCI) have been made to decode phonemes, sub-words, words, or sentences using invasive measurements, such as the electrocorticogram (ECoG), during auditory speech perception, overt speech, or imagined (covert) speech. Decoding sentences from covert speech is a challenging task. Sixteen epilepsy patients with intracranially implanted electrodes participated in this study, and ECoGs were recorded during overt speech and covert speech of eight Japanese sentences, each consisting of three tokens. In particular, Transformer neural network model was applied to decode text sentences from covert speech, which was trained using ECoGs obtained during overt speech. We first examined the proposed Transformer model using the same task for training and testing, and then evaluated the model’s performance when trained with overt task for decoding covert speech. The Transformer model trained on covert speech achieved an average token error rate (TER) of 46.6% for decoding covert speech, whereas the model trained on overt speech achieved a TER of 46.3% (*p >* 0.05; *d* = 0.07). Therefore, the challenge of collecting training data for covert speech can be addressed using overt speech. The performance of covert speech can improve by employing several overt speeches.

## Introduction

Brain–computer interfacing (BCI) is a technique that allows direct communication and control between the brain and external devices through neural signal interpretation^1,2^. BCI has various applications, from assisting individuals with severe physical disabilities to enhancing human–computer interaction and cognitive augmentation. The applications include wheelchair control^3^, cursor control^4^, and rehabilitation^5^.

Speech BCI, which enables speech by decoding human thoughts, is expected to be used by aphasic patients and as a new communication tool in the future^6^. Several techniques to achieve BCI using an invasive electrocorticogram (ECoG) measured by electrodes implanted in the skull are under development. The ECoG is superior to the surface electroencephalogram (EEG) in terms of its spatial and temporal resolution and signal-to-noise ratio; it is particularly suitable for analyzing brain activity related to speech^7,8^.

It has been demonstrated that phonemes, sub-words, words, or sentences can be decoded based on auditory speech perception, overt speech, or imagined (covert) speech. To enable isolated-word speech decoding, Pei et al. used a naive Bayes classifier for overt and covert speech^9^, and Martin et al. employed a support vector machine for auditory speech perception, overt speech, and covert speech^10^. To enable sentence-based speech decoding, Viterbi decoding with a hidden Markov model was applied by Herff et al. for overt speech^11^ and by Moses et al. for auditory speech perception^12,13^.

Deep learning techniques, such as automatic speech recognition (ASR), can decode sentences from overt speech. Sun et al. used a combination of a long short-term memory (LSTM) recurrent neural network model (RNN)^14^ and a connectionist temporal classification decoder^15^ for overt speech and silently articulated speech^16^. Makin et al. successfully applied a sequence-to-sequence (seq2seq) network composed of an encoder and a decoder with bidirectional LSTM (BLSTM) for overt speech^17^. Moreover, an effective method for training the network with a limited amount of ECoG signals has been proposed wherein the network intermediate layer is trained to output speech-latent features with lower dimensions than the input features^17–19^. Sun et al.^16^ and Makin et al.^17^ trained LSTM layers in a seq2seq encoder using mutually synchronized ECoG signals and Mel frequency cepstral coefficients (MFCCs) as inputs and outputs, respectively. Komeiji et al.^20^ introduced the Transformer, a successful neural network model in the field of natural language processing (NLP)^21,22^, to an encoder of the seq2seq model. It achieved a higher accuracy for spoken sentence decoding than the model with the BLSTM encoder^17^ and revealed that the MFCCs used for training do not have to strictly synchronize with ECoG signals.

Research on covert speech decoding into text sentences is the most direct path toward the realization of speech BCI. Some studies have demonstrated advances in covert speech decoding. Pei et al.^8^ and Martin et al.^10^ reported that some cortex areas and high gamma bands of ECoG signals for covert speech are common for overt speech. Angrick et al.^23^ decoded the synthesized speech of isolated words from covert speech using stereotactic EEG (sEEG) electrodes. They used a linear discriminant analysis model trained with sEEG signals of overt speech. Martin et al.^24^ decoded covert speech sentences in ECoG signals into spectrotemporal features by employing a linear mapping model trained using overt speech. Proix et al.^25^ demonstrated that covert speech can be decoded from low- and high-frequency power and cross-frequency as intracranial EEG features. Therefore, owing to such findings, deep-learning techniques can be applied to decode covert speech and pave the way to realizing the practical speech BCI.

To advance the research on text sentence decoding from covert speech, we posit that employing deep-learning techniques is a viable choice. However, collecting covert speech as training data for deep-learning models presents challenges because participants must consciously read sentences silently while seated on special equipment.

To address this issue, we hypothesize that ECoG signals of overt speech can be used as training data for sentence decoding of covert speech. This hypothesis is inspired by previous studies by Oppenheim et al.^26^, which suggested that speech and covert speech production slightly share a common neural substrate. Thus, to validate our hypothesis, we recorded the ECoG signals of two types of speech: overt and covert speech with the same Japanese sentences. To facilitate the recording of the overt and covert speech, we utilized a “karaoke-like text highlighting” technique that aided participants in reading the text aloud or silently. The recorded ECoG signals were used to train seq2seq networks of the Transformer or BLSTM networks that decoded text sentences from ECoG signals. Moreover, we examined the brain regions contributing to sentence decoding to analyze the factors that enable the proposed method to estimate sentences in different tasks.

## Results

In this study, ECoG and audio signals were recorded from 16 volunteer participants (js1 to js16) while they performed two tasks, imagining, or speaking Japanese sentences (i.e., a “covert” task or an “overt” task, as shown in Fig. 1a). The sentences comprised three Japanese phrases, each with two options. This resulted in eight sentences (Fig. 1b). The decoding pipeline to decode text from the ECoG signals denoted as “overt” and “covert” speech, which were recorded during the two tasks, comprises the feature extraction and neural network. This network includes the temporal convolution and the seq2seq network model: the encoder and decoder (Fig.1c). In this study, we implemented the BLSTM or Transformer to the seq2seq model. We trained this model for each participant and each speech (i.e., “overt” or “covert”). In addition, a “shuffle model” was trained on the temporally shuffled ECoG signal to determine if the decoding from the ECoG signal by the Transformer was successful by chance. In the training process, audio signals recorded alongside the ECoG signal were utilized to train the encoder to generate their corresponding MFCCs. Specifically, for training the “covert model,” the MFCCs used were derived from audio signals during the “perception” task, as no audio signals were available during the “covert” task. For more details on the participants, data acquisitions, experimental design, sentences, decoding pipeline, ECoG and audio preprocessing, neural network architecture, and neural network training, see the Methods section.

**Figure 1.**
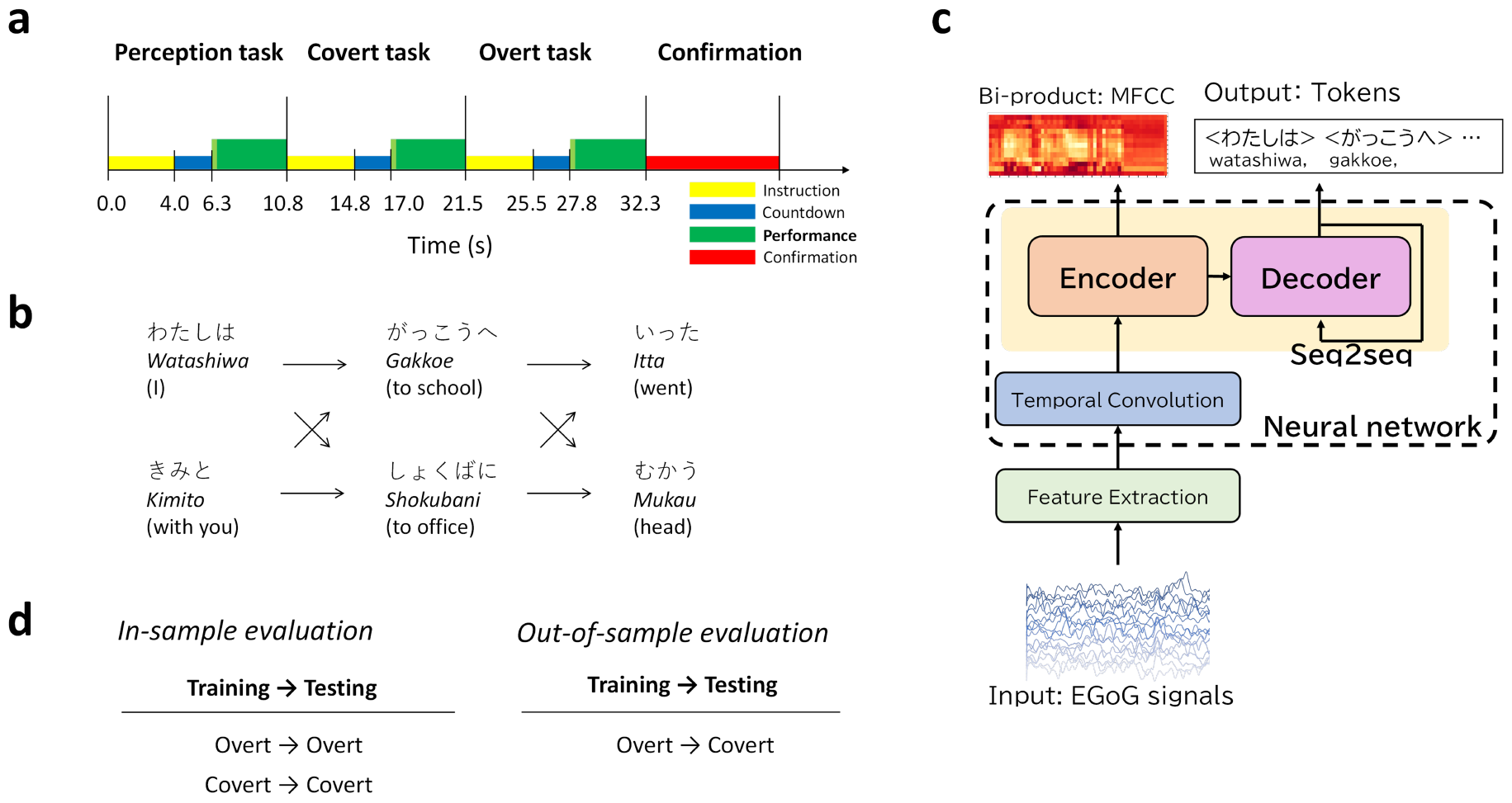
Study overview. **a** Experimental design. A single track includes over 33 s to record sentences for the three tasks. The yellow, blue, and green boxes indicate task instructions, countdown for preparation, and task performance, respectively. The red box is the prompt status for error judgment. **b** Sentence generation. Sentences comprising three Japanese phrases, each with two options, were used. This resulted in a total of eight sentences. The figure depicts the structure of each sentence, with Hiragana representation at the top, Romaji (pronunciation) in the middle, and English translation in parentheses below. **c** The decoding pipeline. The pipeline comprises the feature extraction and neural network, which comprises the temporal convolution, seq2seq network model’s encoder, and decoder. ECoG signals are input into the feature extraction. Token sequences as sentences are output from the decoder. **d** In-sample evaluations showing that the two speeches for training and testing are the same, including overt and covert. Out-of-sample evaluation shows that the two speeches for training and testing are different, including “Overt → Covert.”

We evaluated the decoding performance using the average token error rate (TER) for all decoded sentences. The average TER was computed based on 800 resulting sentences per participant. Subsequently, we utilized these average TERs to evaluate two models: the “overt model” and “covert model” for text decoding from two types of speech: “overt speech” and “covert speech.” Our evaluations included two in-sample evaluations (experiment 1) and an out-of-sample evaluation (experiment 2), which are shown in Fig.1d. For in-sample evaluations, both training and testing utilized the same type of speech, such as overt speech for “Overt→Overt” and covert speech for “Covert →Covert.” In contrast, out-of-sample evaluation involved different speech types for training and testing. Specifically, “Overt→Covert” used overt speech for training but covert speech for testing. Notably, the comparison between the “Overt→Covert” and “Covert→Covert” validated our hypothesis: ECoG signals of overt speech can be used as training data for the sentence decoder of covert speech. In this study, including the validations for our hypothesis, we conducted six comparisons between two results for decoding performance. All the comparisons were assessed using a one-sided Wilcoxon signed-rank test, and the resulting *p*-values underwent Holm–Bonferroni correction for the six comparisons. The number of participants was considered in all comparisons; *n* = 16. Additionally, the effect sizes for ten comparisons were computed using Cohen’s *d*. See the evaluation method and items in the Methods section for more details.

### Decoding Performance

The experiments yielded three findings. First, the Transformer model outperformed the BLSTM model. Second, the Transformer models trained on covert and overt speech exhibited significantly lower TER in text decoding from covert speech compared to “shuffle models.” This indicates that the Transformer model can extract meaningful information from covert speech. Third, the performance using the “overt model” was comparable to that of the “covert model,” regarding text decoding from covert speech using Transformer models. This finding supports our hypothesis that ECoG signals of overt speech can be used as training data for the sentence decoder of covert speech.

The TERs for all 16 participants are plotted in Fig. 2 as dots representing each participant’s TER and box plots. Figure 2a shows the results of experiment 1: in-sample evaluation and Fig. 2b shows the results of experiment 2: out-of-sample evaluation. Each figure includes the comparison of the BLSTM and Transformer models. Additionally, the results of the six comparisons depicted in the figure based on *p*-values and effect sizes are listed in Suppl. Table 1.

**Figure 2.**
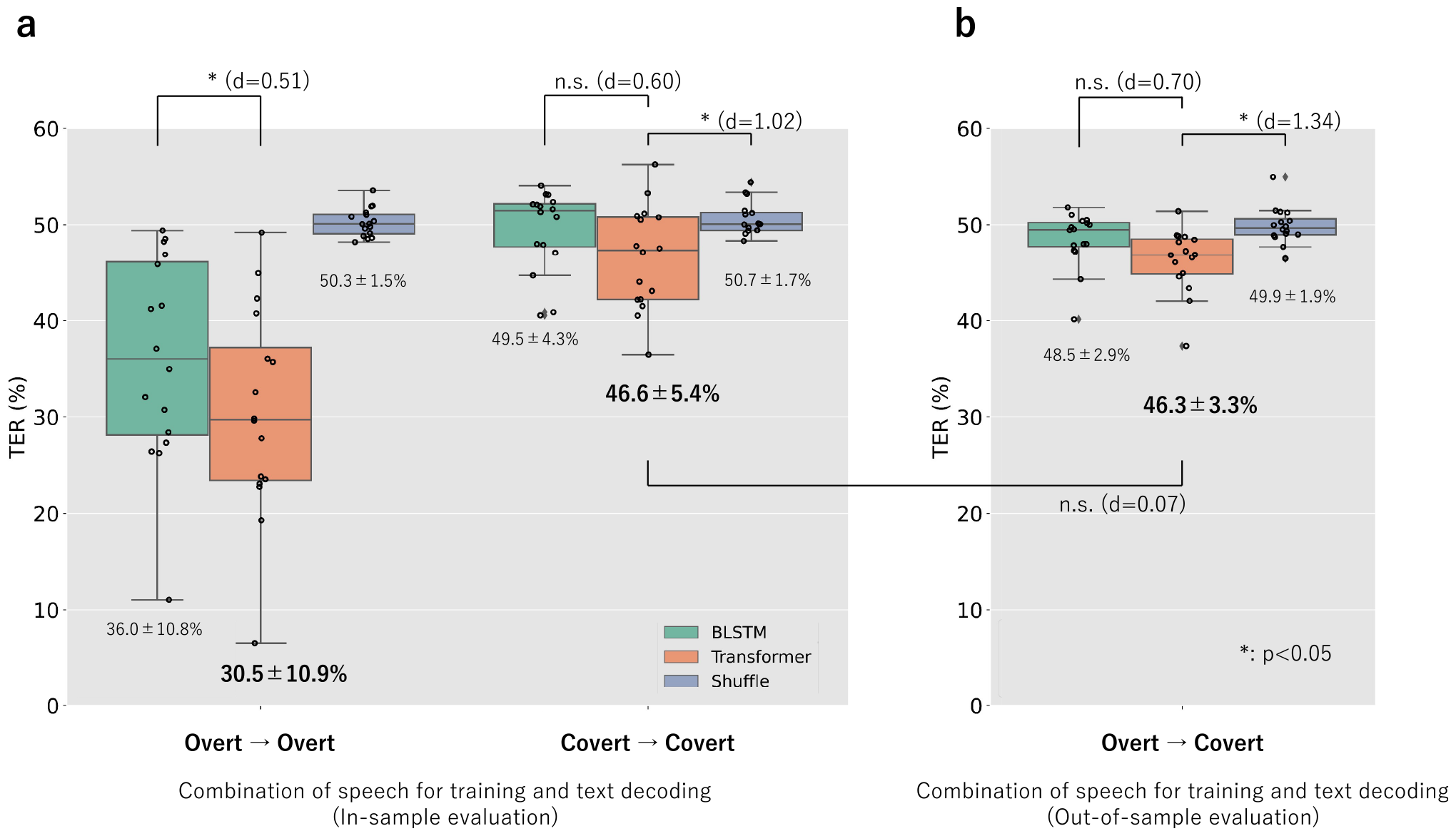
Decoding performance. Boxplot depicting the distribution of the token error rate (TER) for five combinations of speech for training and testing. Each dot on the plot represents an individual participant. The green boxes represent the TER distribution of the BLSTM seq2seq model, whereas the orange ones represent the Transformer seq2seq model. The blue ones show the Transformer seq2seq models labeled as “Shuffle,” which were trained with the temporally shuffled ECoG signals. We defined outliers as values that are 1.5 times the interquartile range (IQR) away from the median. The IQR represents the range of values within the middle 50% of the data. The significance (∗: *p <* 0.05, n.s. : *p* ≥ 0.05) was calculated with the one-side Wilcoxon signed-rank test Holm—Bonferroni corrected for ten comparisons. For each comparison, the effect size using Cohen’s *d* is included. The numbers in the figure indicate the average ± standard deviation. **a** In-sample evaluation. **b** Out-of-sample evaluation.

With respect to the first finding, the Transformer model outperformed the BLSTM in all cases. For the cases of “Overt→Overt,” the Transformer significantly outperformed the BLSTM, with small *p*-value of 3.9*×*10^−4^. The effect size, quantified by Cohen’s *d*, was medium to large, with values of 0.51. For “Covert→Covert” and “Overt→Covert,” although the Transformer did not exhibit a significant advantage over the BLSTM (*p*-values of 6.8*×*10^−2^ and 0.11), the effect sizes remained medium at 0.60 and 0.70. For the second finding, the Transformer model showed significantly superior decoding performance for covert speech in both “Covert→Covert” and “Overt→ Covert” compared to the “shuffle model” (*p*-values of 3.0*×*10^−2^ and 1.4*×*10^−3^). Moreover, the effect sizes were as large as 1.02 and 1.34. For the third finding, regarding the decoding of covert speech using Transformer models, the performance using the “overt model” (TER = 46.3%) was comparable to that using the “covert model” (TER = 46.6%). The *p*-value was as large as 2.9 and the difference in performance was as small as an effect size of 0.07.

### Anatomical analysis

We investigated the cortical areas contributing to text decoding for both model training and text decoding for each speech. The contribution was determined by generating saliency maps representing the text output’s gradients to the input ECoG segments. Figure 3 illustrates the electrode contributions projected onto the 3D cortex object, as demonstrated by one example participant, js11. Participant js11 achieves lower TERs than the median among all participants for three combinations of speech for training and testing. The model that achieved lower TERs is expected to generate more reliable saliency maps. Details regarding electrode contributions from the remaining participants are provided in the supplementary material. The results in Fig. 3 indicate that the trend of electrode contributions is primarily influenced by the differences among the models used rather than the speeches for decoding. This trend is consistent among all the participants (see supplementary material). For participant js11, when the “overt models” are used for text decoding, the sensorimotor cortex (SMC) exhibits the strongest contribution, regardless of the decoding targets. This observation extends to five out of eight participants with TERs below the median in the “Overt→Overt,” where SMC emerges as the predominant contributing area. For the “covert models,” the contributed area is less prominent than that of the “overt models.” Consistent contributions from specific brain areas across different models, regardless of model type (e.g., SMC for the “overt model”), include participants js1 and js14 (see Supple. Fig. 4 and Fig. 17). For js1, the main contributing area was the inferior parietal cortex, whereas for js14, it was the occipital lobe.

## Discussion

The results herein suggest that a machine-learning model trained by overt speech is applicable for decoding covert speech. This section discusses the results from four aspects. First, the BLSTM and Transformer models applied to the encoder and decoder parts of the seq2seq network are compared. Second, the effectiveness of MFCCs in training the encoder part of the network is examined. Third, we discuss the main topic of this study. Both the “covert model” and “overt model” can decode text from covert speech. We explain this finding by analyzing the distribution of electrode contribution shown in the saliency maps. Finally, we discuss the prospects for realizing speech BCI based on the results of this study.

First, this study compared BLSTM and Transformer models for text decoding from ECoG signals. While some studies showed that the Transformer model outperformed LSTM or BLSTM models in ASR^27,28,29^, the studies on EEG signals have also successfully applied Transformer models for BCI^30,31,32,33,34,35,36^. Notably, studies by Tao et al.^31^ and Siddhad et al.^32^ showed that Transformer models outperformed BLSTM models, even when they were trained on a small dataset, that is, of less than 10 h. This dataset size is considerably smaller than that typically employed in ASR, which usually comprises over 1,000 h of speech. Moreover, Tao et al.^31^ demonstrated the effectiveness of their gated Transformer approach using an image classification task for a human brain-visual dataset and a motor imagery classification task. The brain-visual dataset^37^ was collected from six participants and contains 11,964 EEG segments of 0.5 s each, whereas the motor imagery dataset^38^ was collected from 109 participants and contains 11,345 EEG segments of 4 s each. Siddhad et al.^32^ explored the efficacy of the Transformer model for the classification of raw EEG data. The performance of the Transformer model was evaluated on an “Age and Gender Dataset”^39^ and a simultaneous task EEG workload (STEW) dataset^40^. The former dataset consists of raw EEG signals from 60 participants. The data were acquired for ten sessions of 10 s for each participant. The latter dataset consists of raw EEG signals from 48 participants. For each participant, the data were acquired for 5 min.

Komeiji et al.^20^ demonstrated the efficacy of the Transformer using overt speech with eight participants. Each participant-dependent model for text decoding was trained on a significantly small amount of data (approximately 3 min), consisting of 64 segments (approximately 3 s each). Our study extended this approach by examining covert speech. The Transformer outperformed BLSTM even with small datasets for training, as demonstrated in the above studies. Specifically, our results indicate that the Transformer’s ability to capture long-term dependencies through a self-attention mechanism, which encodes information from the entire sequence, is crucial in its superior performance for all three types of speech in ECoG signals. These findings are significant as they suggest that the Transformer will remain a dominant approach in brain activity modeling, especially when managing limited training datasets, paving the way for future research in BCI.

In the second aspect, we investigated the effectiveness of using MFCCs in training the encoder part of our model. Prior studies have proposed an effective method for training a network with a limited number of ECoG signals, where the intermediate layer of the network is trained to output speech-latent representations with lower dimensions than the input features^18,16,19,17,20^. While Makin et al.^17^ required synchronization of speech for calculating MFCC and ECoG signals, Komeiji et al.^20^ discovered that synchronization was unnecessary. In this study, we applied MFCC to train our models using ECoG signals during covert speech. Suppl. Fig. 3 shows that despite their difficulty in synchronizing the covert speech, MFCCs play an important role in training the models. This is because the self-attention mechanism of the Transformer, which allows it to encode information from the entire sequence, absorbs the asynchrony between inputs and outputs. Future work will include a detailed analysis of the output of the intermediate layers, such as the attention layers.

Although the MFCCs used to train the encoder play an underappreciated role, as they are discarded during the decoding phase, the output of the encoder can be utilized for speech synthesis. Instead of MFCCs, Shigemi et al.^41^ demonstrated that the Transformer encoder can generate Mel spectrograms as outputs when the overt speech is used as the input. These Mel spectrograms can be fed into a neural vocoder called Parallel WaveGAN^42^ to synthesize high-quality audio speech. It is worth considering the integration of synthesized speech into an already-trained ASR system, which is an alternative approach to the seq2seq model composed of an encoder and a decoder proposed herein.

In the third aspect, we explore why the “overt model” performs similarly to the “covert model” in text decoding from covert speech. We refer to the saliency maps in Fig. 3 and previous studies to analyze the electrode contribution distribution. The electrode contribution distributions of the “covert model” were broader than those of the “overt model” for most participants (14 out of 16). According to Tian et al.^43^, the cortical activity of covert speech involves a broader area in the cortex. The authors found that articulation imagery induced greater activity in frontal-parietal sensorimotor systems, including the sensorimotor cortex, subcentral (BA 43), middle frontal cortex (BA 46), and parietal operculum (PO). The “covert model” can be trained to focus on these areas, resulting in a broader electrode contribution distribution.

**Figure 3.**
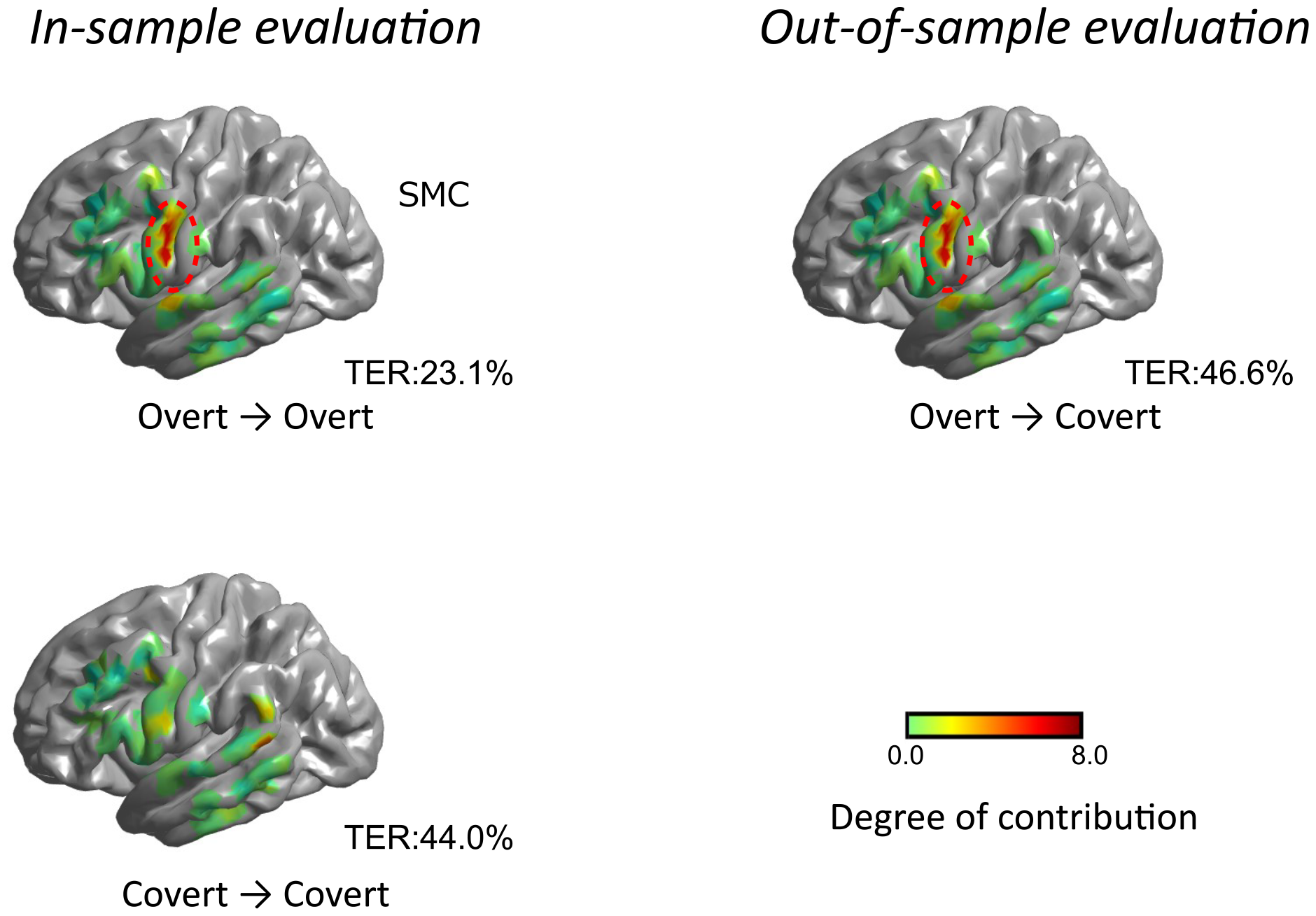
Electrode contributions for text decoding on averaged FreeSurfer pial surface image. The electrode contributions are illustrated using a color map, where green and red indicate the weak and strong contributions, respectively. The left column indicates the results of the in-sample-evaluation: “Overt → Overt” and “Covert → Covert.” The right column indicates the results of the out-sample-evaluation: “Overt → Covert.” The average TERs in js11 are aligned below the cortex objects. The cortex objects are marked with red dashed circles to emphasize the areas that make significant contributions to text decoding. In the case of the “overt model,” the sensorimotor cortex (SMC) is the highlighted area.

In contrast, for the “overt model,” certain SMC electrodes contributed strongly to text decoding for eight participants. It has been shown that the activities of SMC contribute to decoding text from overt speech^8,10,17^. The results of this study are consistent with the results of these studies. “Overt models” can be trained to focus on strong activities related to speaking in specific brain areas such as the SMC. “Overt models” can decode text from covert speech, suggesting that there may be common signal patterns in those brain areas for overt and covert speech.

Although this “commonality” of the activities for overt and covert speech has been introduced in prior studies^44,25^, how the activities are generated remains unclear. There are two hypotheses in covert speech production: the motor hypothesis and abstraction hypothesis^25^. The motor hypothesis argues that covert speech is an attenuated version of overt speech^25,45,26,46^. Oppenheim et al. found evidence that covert speech might be impoverished at lower (featural) levels, but its production at higher (phonemic) levels is unclear^45^. The abstraction hypothesis argues that covert speech originates from higher-level linguistic representations. Price et al. argue that articulation planning is similar in overt and covert speech^47^. Further analysis of the results could help determine the correct hypothesis, although the current analysis alone is insufficient to conclude.

For two exceptional participants (js1 and js14) out of the 16 participants, the distributions of the electrode contributions across the two speeches (overt speech and covert speech) were nearly identical. In the case of participant js1 (Suppl. Fig. 4), the inferior parietal cortex, which is associated with memory-related activity^48^, showed a significant contribution. For participant js14 (Suppl. Fig. 17), the occipital lobe exhibited notable contributions. The previous study^49^, reported that this area is related to both listening and reading comprehension-related activity. These findings suggest that the activities related to memory and comprehension may be dependent on the two tasks. The electrode contribution distributions observed in js1 and js14 can be considered valid results.

Finally, we discuss the contribution of the results obtained in this study. The result that the “overt model” can be used for text decoding from covert speech may pave the way to realizing practical speech BCI based on machine learning. Although the machine-learning model requires multiple training data, it is difficult to collect the covert speech because the participants must read the “Karaoke-like text highlighting” to obtain labels. In contrast, collecting overt speech is easier as it can be obtained during normal activities, such as daily conversation. Additionally, the audio signal accompanying the speaking task can be used as label data for training. If more overt speech is collected, the accuracy of text decoding from covert speech may significantly improve.

## Conclusion

This study demonstrated that a machine learning model trained on ECoG signals recorded during overt speech can decode text sentences from covert speech. ECoG signals for overt and covert tasks, related to eight Japanese sentences consisting of three tokens, were recorded from 16 experimental participants with electrodes implanted intracranially for epilepsy treatment.

The decoding performance using the model trained by the overt speech (TER = 46.3%) was almost the same as that using the model trained by the covert speech (TER = 46.6%; the *p*-value was more than 0.05 as derived via the one-side Wilcoxon signed-rank test, and Cohen’s *d* was 0.07). The anatomical analysis showed the possibility of the common signal patterns of activities in certain cortex areas for covert and overt speech. Further analysis using the ECoG signals recorded in this study on common signal patterns in covert and overt speech may help elucidate how speech-related activities are generated. Because the “overt model” can be applied in text decoding from covert speech; thus, the collection of training data is much easier and may accelerate the progress of speech BCI.

## Methods

### Participants

The 16 volunteer participants in this study (js1 to js16; according to Table 1, eight males, eight females, mean age 25.2 years, range 8–41) were under treatment for drug-resistant temporal lobe epilepsy at the Department of Neurosurgery, the Juntendo University Hospital. ECoG electrode arrays were surgically implanted on each participant’s cortical surface to localize their seizure foci. ECoG and audio data in this study were obtained from all 16 human participants who gave informed consent. The experimental design to record the data was approved by the Medical Research Ethics Committee of Juntendo University Hospital. The data analysis was approved by the Ethics Committee of the Tokyo University of Agriculture and Technology. All research was performed in accordance with relevant guidelines and regulations.

**Table 1.**
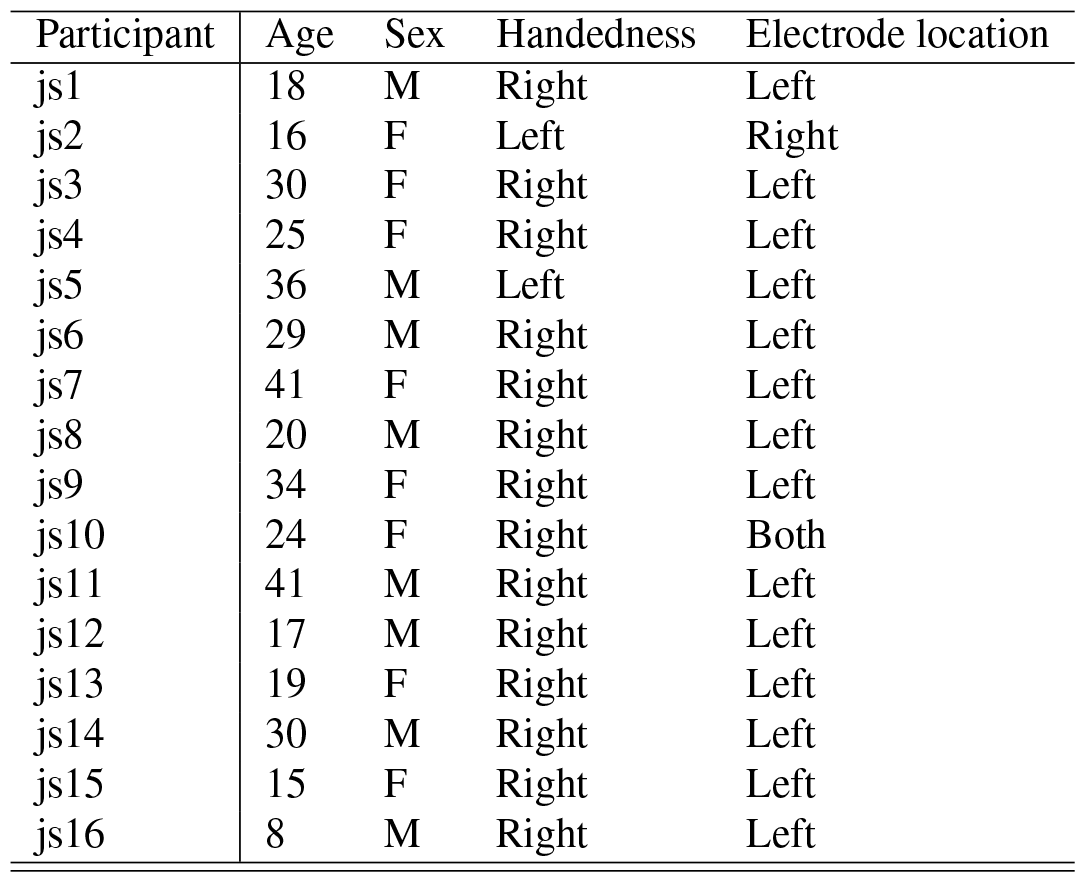
Clinical profiles of the 16 participants. The age, sex, handedness, and electrode location are listed. The sampling rates were 1,200 Hz for three participants (js1 to js3) and 9,600 Hz for other participants (js4 to js16).

| Participant | Age | Sex | Handedness | Electrode location |
| --- | --- | --- | --- | --- |
| js1 | 18 | M | Right | Left |
| js2 | 16 | F | Left | Right |
| js3 | 30 | F | Right | Left |
| js4 | 25 | F | Right | Left |
| js5 | 36 | M | Left | Left |
| js6 | 29 | M | Right | Left |
| js7 | 41 | F | Right | Left |
| js8 | 20 | M | Right | Left |
| js9 | 34 | F | Right | Left |
| js10 | 24 | F | Right | Both |
| js11 | 41 | M | Right | Left |
| js12 | 17 | M | Right | Left |
| js13 | 19 | F | Right | Left |
| js14 | 30 | M | Right | Left |
| js15 | 15 | F | Right | Left |
| js16 | 8 | M | Right | Left |

### Data acquisition

The ECoG signals were recorded using intracranial electrodes implanted on the cortical surface. We set the ground (GND) as an epidural electrode. All signals other than the one with the epidural electrode, between their respective electrodes and the GND as a reference, were amplified using a biomedical amplifier (g.HIAMP, g.tec Medical Engineering GmbH, Austria). The electrooculograms were recorded simultaneously with the ECoG signals by employing four disposable electrodes to confirm eye movements. In addition to ECoG signals, audio signals, such as participants’ utterances, were recorded using a microphone (ECM360, SONY, Japan) and digitized in sync with the ECoG signals.

The ECoG and audio signals were digitized at sampling rates of 1,200 Hz for three participants (js1 to js3) and 9,600 Hz for other participants (js4 to js16) (The use of the low sampling rate of 1,200 Hz for participants js1 to js3 was an accident during the experiment. However, we did not discard them from the analysis).

Using a photodetector circuit, we recorded trigger signals indicating changes in the event status. The circuit combined a linear light sensor (LLS05-A, Nanyang Senba Optical and Electronic Co., Ltd., China) with an operational amplifier (NJM3404AD, New Japan Radio, Japan). The photodetector was placed in a monitor corner where the pixel colors changed in response to the event status, capturing and recording the changes. We used the Simulink toolbox (MathWorks, USA) to control the biomedical amplifier and record all observed signals.

### Experimental design

To record the three conditions of ECoG signals (i.e., auditory speech perception, overt speech, and covert speech), we designed the task schedule depicted in Fig. 1a. According to the task schedule, participants performed perception, overt, and covert tasks. We refer to the series of three tasks as a “track.” A single sentence, which was presented at random, was recorded for each track. More than 6 s intervals between tasks can prevent time-correlated artifacts. Participants repeated this track 80 times (if they made a mistake, it would take more than 80 times), and we obtained 80 segments of ECoG and audio signals for each task and participant. During the recording, the participants were seated on a chair with a desk equipped with a screen, loudspeaker, microphone, and keyboard. The monitor and loudspeaker were operated based on the task schedule.

The track consisted of the perception, overt, and covert tasks in that order. At the beginning of each task, the instruction and countdown periods were inserted to prepare the participants for the task (yellow and blue boxes depicted in Fig. 1a). In the perception task, the participants were asked to listen to the sentence played from the loudspeaker and gaze at the cross symbol displayed on the monitor. In the covert and overt tasks, the participants were asked to read silently or aloud a sentence characterized as “Karaoke-like text highlighting” displayed on the monitor. The speed of text highlighting was at a constant rate (5 characters/s). To avoid reading faster or slower than the highlight speed at the beginning of a sentence, we inserted a dummy word, which does not exist in the Japanese words and has no meaning but can only be pronounced, at the beginning of the sentence (lime green boxes in Fig. 1a). At the end of the track, the participants operated the keyboard to judge whether this track had any mistakes (red box in Fig. 1a). The next track started if the participant judged that there was no mistake; otherwise, this track was repeated.

During the experiments, we continuously recorded ECoG signals, an audio signal, and a trigger signal, which indicated the periods of the dummy words and sentences (lime green and green boxes in Fig. 1a). The trigger signal was used for trimming both the ECoG and audio signals.

### Sentences

The sentences displayed to the participants comprised three Japanese phrases (Fig. 1b). The total number of sentences was eight. Thus, the same sentence appeared ten times for all 80 trials of each task. Each phrase had two candidates to generate one sentence. The first phrase was either “watashiwa” (I) or “kimito” (with you), the second phrase was either “gakko:e” (to school) or “shokubani” (to the office), and the third phrase was either “itta” (went) or “mukau” (head). Consequently, we generated eight (= 2 *×* 2 *×* 2) sentence patterns.

### Tokenization of phrases

To apply six phrases composing eight sentences and the two additional phrases “start of sentence” and “end of sentence” to the outputs of a neural network model, we assigned each phrase a token ID consisting of an integer value from zero to seven. The number assignment to each phrase is as follows:

- “watashiwa” (I) : 0
- “kimito” (with you): 1
- “gakko:e” (to school): 2
- “shokubani” (to the office): 3
- “itta” (went): 4
- “mukau” (head): 5
- start of sentence: 6
- end of sentence: 7

### Decoding pipeline

We decoded the sentences in text form from each segment of the ECoG signals by employing the decoding pipeline shown in Fig. 1c. The pipeline comprises the feature extraction and neural network parts, which comprise the temporal convolution, seq2seq network model’s encoder, and decoder.

In the feature extraction part, the segments of the ECoG signals were converted to envelopes of high gamma bands with a sampling rate of 200 Hz. A detailed explanation of the feature extraction part is presented in Section ECoG preprocessing. The envelopes were input to the temporal convolution part to downsample the time axis effectively. The output of the temporal convolution was input to the seq2seq network. The encoder part of the seq2seq network was characterized to be trained to output the MFCC extracted from the segment of the audio signal, which was paired with the segment of the ECoG signal. Detailed explanations of the calculation of the MFCC and the training of the seq2seq network are provided in Sections Audio preprocessing and Neural network training. In text decoding, the encoder part outputs the estimated MFCC as a bi-product, and we obtained the output text from the decoder part.

In this study, we implemented the seq2seq networks as two types, BLSTM-based and Transformer-based. In the BLSTM-based seq2seq, we applied the BLSTM to the encoder and the LSTM to the decoder. This architecture is the same as that used by Makin et al.^17^. In the Transformer-based seq2seq, we applied the Transformer to both the encoder and decoder. The Neural network architecture section provides a detailed explanation of the architecture.

### ECoG preprocessing

Preprocessing, which obtains the ECoG features from the ECoG signals used for training and decoding in a machine learning network, entails the following steps. In the first step, electrodes with artifacts and excessive noise were removed by visual inspection. Consequently, the numbers of electrodes used for the participants in the analysis were as follows: 72 for js1, 48 for js2, 48 for js3, 54 for js4, 54 for js5, 54 for js6, 53 for js7, 55 for js8, 71 for js9, 42 for js10, 45 for js11, 39 for js12, 56 for js13, 58 for js14, 65 for js15, and 24 for js16. The valid electrode coverage across all participants is depicted in Suppl. Fig. 1. In the second step, we trimmed each segment from both the ECoG and audio signals based on the start and end times of sentences from the trigger signal; specifically, we discarded the periods of the dummy words. The length of each trimmed segment was the maximum duration of all sentences throughout the experiment for each participant. This operation prevented the text decoder from predicting sentences based on their trimmed length. The trim operation before signal processing, such as filtering, can prevent time-correlated artifacts that are the effects of the previous segments. In the third step, the trimmed ECoG signals were anti-aliased (low-pass filtered at 200 Hz) and downsampled to 400 Hz. IIR notch filters of 50 and 100 Hz were applied to suppress the power supply noise and the harmonic. Finally, the amplitudes of the analytic signals at each electrode were extracted in each band of eight adjacent frequency bands, between 70 and 150 Hz, filtered by eight finite impulse response (FIR) bandpass filters, averaged across bands, and downsampled to 200 Hz. The eight FIR bandpass filters were designed with passbands of 68–78, 74–84, 82–92, 91–102, 101–112, 112–124, 124–136, and 137–150 Hz. The amplitudes of the analytic signals were then z-scored, and the ECoG features were obtained.

### Audio preprocessing

The trimmed audio signals were converted to MFCCs using the “Python speech features package^50^.” The calculation parameters were set to 20 ms as the window length and 5 ms as the window step. For the training of the “covert model,” MFCCs were obtained from the audio speech during the perception task, while for the training of the “overt model,” they were computed from the audio speech during the overt task. The audio signals were discarded for js1, js2, and js3 because their sampling rate was 1,200 Hz, with incomplete speech information. Therefore, throughout the experiment, the MFCCs for js1, js2, and js3 were computed from an audio signal of js4 instead. The asynchrony caused by substituting the audio speech during the overt task was inevitable for these three participants.

### Neural network architecture

We applied the BLSTM and Transformer to the seq2seq model for text decoding. BLSTM^14^ is a type of RNN explicitly designed to handle both long-term and short-term memory efficiently. It finds common applications in ASR and NLP. However, the Transformer^21^ is a neural network model distinguished by its attention layers, and is well-suited for capturing long-term dependencies. Since 2018, the Transformer has emerged as the de facto standard in machine learning technology, in ASR and NLP as well as in diverse fields, including image processing.

Here, we explain the detailed network architecture based on the Transformer-based seq2seq model. The network architecture is depicted in Fig. 4, and the implementation parameters are described in Table 2. The network processes the sequences in three stages: a temporal convolution network, an encoder network, and a decoder network. As shown in Fig. 4, the architecture is characterized by the use of the Transformer for both the encoder and decoder stages, and the encoder is characterized to output MFCCs.

**Figure 4.**
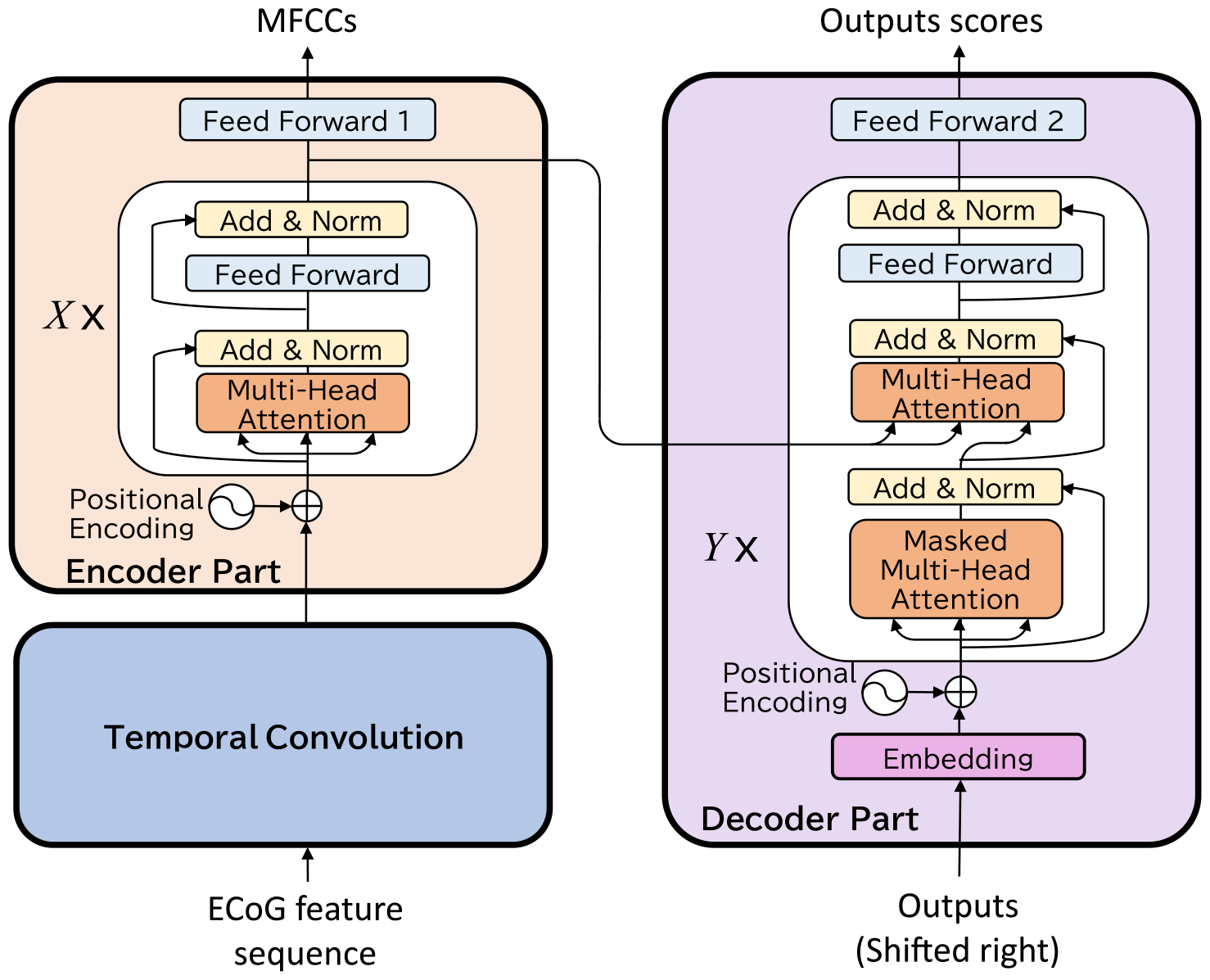
Detailed neural network architecture.

**Table 2.**
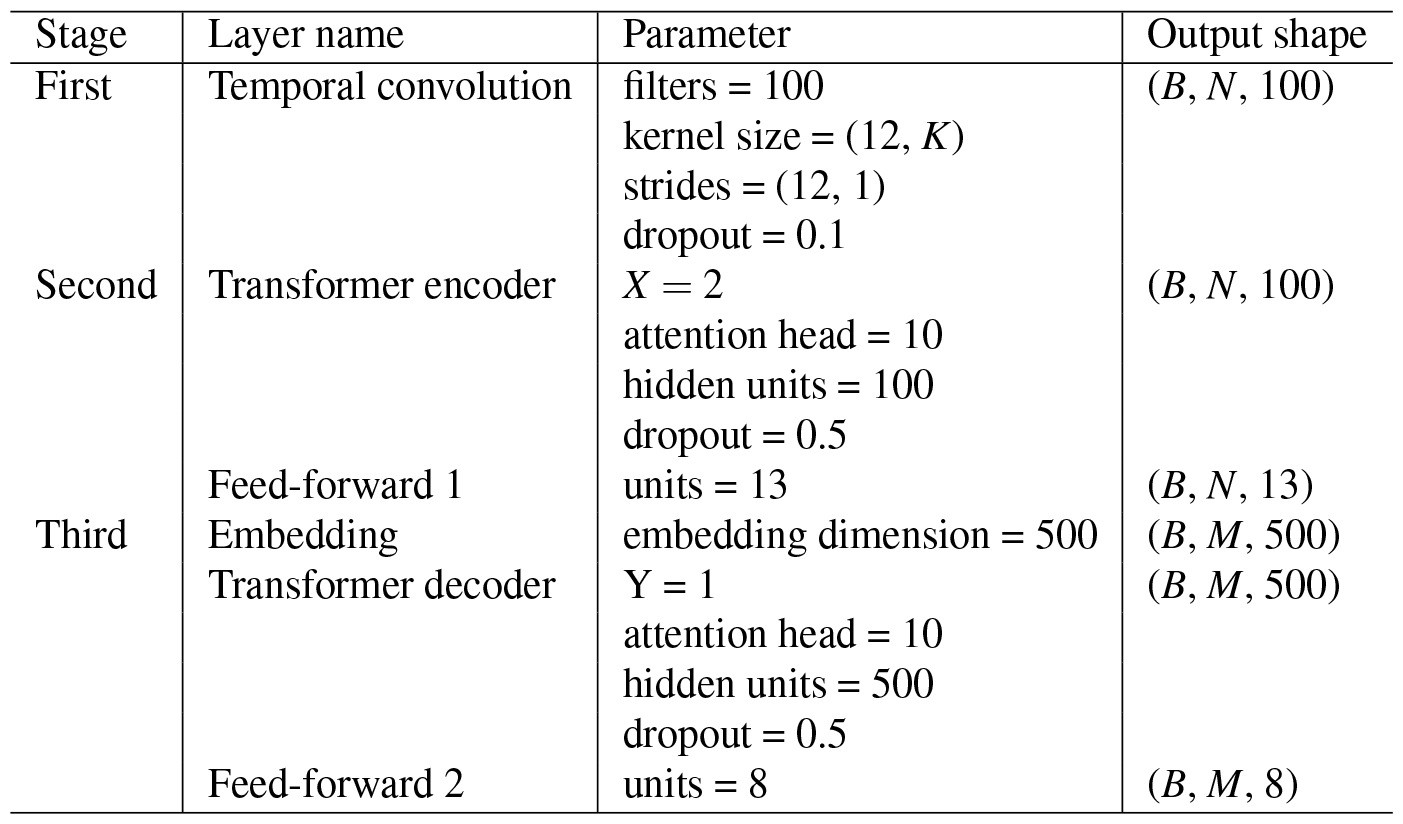
Network parameters.

In the first stage, the temporal convolution (convolutional neural network; CNN) layer effectively downsampled the ECoG features. The input to the CNN consisted of *K* features, each of length *L* (where *K* represents the number of electrodes). The number of electrodes *K* ranged from 24 to 72 among all 16 participants, and the length *L* was approximately 700 (3.5 s sequence of a 200 Hz feature). Each convolutional filter with a kernel size of *W×K* was applied to the *K* features with a stride of *W* ; consequently, the CNN layer output a sequence of *N* (= ceil(*L/W* )) vectors of *C* with a dimension of *C*.

In the second stage, the Transformer encoder received the outputs of the temporal convolution layer and comprised a stack of *X* identical layers, each with two sub-layers. The first sub-layer was the multi-head self-attention mechanism, and the second was a fully connected feed-forward network. Furthermore, these two layers had a residual connection, and layer normalization was applied to their outputs. The outputs of the Transformer encoder were the sequences of feature vectors of dimension *C* with a length of *N*, the same size as the inputs. The outputs of the stack of *X* identical layers were input to the decoder layer and the feed-forward network called “feed-forward 1” to represent the sequence of MFCCs. The “feed-forward 1” layer played an important role in regularizing the outputs of the Transformer encoder. These outputs spanned a latent feature space with lower dimensions and mitigated the problem of limited training data. During training, the Transformer encoder was trained to approximate the outputs to the sequence of MFCCs whose length was the same as that of the outputs of the temporal convolution layer *N*. For example, when the 200 Hz feature is input into the temporal convolution layer with a stride of *W* = 12, the time step of the temporal convolution layer’s output is 600 ms. MFCCs calculated with a window step of 50 ms should be downsampled with the same stride value of *W* = 12. Notably, the MFCCs were not applied to the encoder output in the decoding phase.

In the third stage, the Transformer decoder received the output of the stack of *X* identical layers in the Transformer encoder. The Transformer decoder was composed of a token embedding layer, stack of *Y* identical layers, and feed-forward network called “feed-forward 2.” The embedding layer transforms the token IDs into the embedding vectors. The *Y* identical layers were similar to the *X* identical layers of the Transformer encoder, except for two differences. The first difference was that the identical layers had an additional masked multi-head attention layer in front of the multi-head attention layer. The second difference was that the multi-head attention layer received the output of the Transformer encoder as the key and query of the attention layer. The embedding layer transformed the token IDs into the embedding vectors. The stack of *Y* identical layers received the *M* length of the embedded vectors and output the *M* length of the vectors. The sequence of the vectors was applied in the “feed-forward 2” to represent the sequence for scores of tokens.

### Neural network training

We trained the neural network based on the parameters described in Table 3. Two types of loss values were introduced to train the networks. The first was the categorical cross-entropy loss between a reference sentence (sequence of text token IDs) and the decoder output sequence of the vectors whose elements represent the token IDs. The second one was the mean square error (MSE) between the reference MFCCs and outputs of the encoder part. These values were linearly combined by the MFCC-penalty weight *λ* of the MSE loss. In this study, we set the weight to be a value of 0.1. The training parameters are listed in Table 3. Some of these parameters were the same as those used by Makin et al.^17^.

**Table 3.** Training parameters.

| Parameter name | Value |
| --- | --- |
| Learning rate | 0.0005 |
| MFCC-penalty weight, $\lambda$ | 0.1 |
| Batch size | 16 |
| # training epochs | 800 |

We trained two types of neural network models: “overt model” and “covert model.” In training the “overt model,” we used pairs of segments of the ECoG for overt speech and the audio signals simultaneously recorded during the overt tasks. In training the “covert model,” we used pairs of the segment of the ECoG signals for covert speech and the audio signal segment during the perception task. Note that the sentence content of the two segments was the same. In training the models for participants js1 to js3, whose audio signals were discarded owing to the low sampling rate recording, we used the audio signals of js4 instead.

### Evaluation method

We evaluated the models using a five-fold cross-validation approach. Training and evaluation in one fold used up to 64 and 16 segments, respectively. In a fold, ten network models were trained and evaluated to prevent variations in the decoding results owing to variations in the initial values of the network weights and variations in the computation results by the GPU. Consequently, up to 800 results (10 models *×* five folds *×* 16 sentences) of a sentence decoding for each speech and each participant were obtained. We evaluated the performance of decoding based on the TER. The TER was calculated by the following equation:

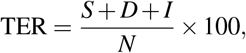

where *S, D*, and *I* represent the number of errors for tokens: substitution, deletion, and insertion, respectively. *N* indicates the total number of tokens in sentences. TERs were calculated as averages across the 800 results for each task and each participant.

### Evaluation items

We evaluated two models (“overt model” and “covert model”) for decoding text from two types of speech (overt speech and covert speech). We used two architectures: BLSTM-based and Transformer-based models. We had two types of speech for training, each corresponding to two different models, and two types of speech for testing. Consequently, there were four combinations of models and input speech for testing, where each model was paired with each type of input speech to obtain the output sentences. We divided the four patterns into two groups: the first group included three patterns where the two speeches were the same, and we called this group an “in-sample evaluation.” The other group included two patterns where the two speeches differed, and we called this group an “out-of-sample evaluation.” To demonstrate that the Transformer-based model had learned meaningful information in ECoG signals, it was compared with the model (“shuffle model”) trained on temporally shuffled signals.

***Experiment 1: in-sample evaluation***.

We conducted all two in-sample evaluations as follows:

- Overt → Overt (using the model trained with “Overt” speech for text decoding from “Overt” speech).
- Covert → Covert (using the model trained with “Covert” speech for text decoding from “Covert” speech).

In the expression of the combinations, “*value1* → *value2*” represents text decoding using the model trained with “value1” speech for decoding text from “value2” speech.

***Experiment 2: out-of-sample evaluation***.

We selected one out-of-sample evaluation out of all two as follows:

- Overt → Covert

Notably, the comparisons between the “Overt → Covert” and “Covert→Covert” validate our hypothesis: ECoG signals of overt speech can be used as training data for the sentence decoder of covert speech.

### Statistical analysis

To compare the performance of the two models, we employed a one-sided Wilcoxon signed-rank test under the hypothesis that “Comparison 2 outperforms Comparison 1.” All the comparisons were conducted considering the number of participants *n* = 16. The resulting *p*-values underwent Holm–Bonferroni correction for the six comparisons conducted in this study. In addition to statistical tests, we calculated effect sizes between the two results as Cohen’s *d*.

### Electrode contributions

The electrode contributions were derived based on saliency maps of the neural network model trained with each condition. The straightforward way to determine the contribution of electrodes is to determine the best model for text decoding among the models trained by various subsets of electrodes. The variation for the subsets of electrodes is enormous, and it is time-consuming to train and evaluate all the models. Preliminary evaluation of training and decoding of a single electrode ECoG signal for js11 of “Overt → Overt” showed that the electrodes with lower TERs tended to approximately coincide with those electrodes that had high contributions as derived by the saliency map (Suppl. Fig. 2).

Therefore, we introduced the saliency map^51^ to determine the electrode contribution. The saliency map is the derivative for the output on input for the following form. The shape of the saliency map was the same as that of the input ECoG features, having the contribution value for output. The value of each element of the saliency map is derived for each time point and each electrode. We were interested in determining the trend of electrode contributions across all trimmed ECoG signals for text decoding in each participant. To this end, we computed the variance along the time axis to observe the saliency trend over the time sequence. Our objective was not to obtain the absolute values for each electrode but to compare the relative values among them. To normalize the contribution values across electrodes for a single trial, we calculated their z-score of variances. Finally, the z-scored values for each trial were averaged for the evaluation data sets of all 800 decoding times. The contribution values, calculated as the average of the z-scores as described above, with the electrode positions for each participant, were transformed into Talairach coordinates, and plotted on the averaged FreeSurfer pial surface image, known as “fsaverage”^52,53,54^.

## Acknowledgements

This work was supported in part by JSPS KAKENHI, 20H00235, and 23H00548. We would like to thank Editage (www.editage.jp) for English language editing.

## Notes

### Competing Interest Statement

The authors have declared no competing interest.

### Summary of Updates

- Mejor revision: the experiment results of perception tasks were removed due to some errors in ECoG signals during the tasks. - Minor revisions: some errors in the study were modified. - Supplemental file updated.

