## Supplemental file for "Feasibility of decoding covert speech in ECoG with a Transformer trained on overt speech"

| Comparison 1 | Comparison 2 | $p$-value (Uncorrected) | $p$-value (Corrected) | Effect size |
|---|---|---|---|---|
| Transformer (“O → O”) | BLSTM (“O → O”) | $3.9 \times 10^{-4}$ | $3.9 \times 10^{-4}$ | 0.51 |
| Transformer (“C → C”) | BLSTM (“C → C”) | $1.7 \times 10^{-2}$ | $6.8 \times 10^{-2}$ | 0.60 |
| Transformer (“O → C”) | BLSTM (“O → C”) | $2.2 \times 10^{-2}$ | $1.1 \times 10^{-1}$ | 0.70 |
| Transformer (“C → C”) | Shuffle (“C → C”) | $1.0 \times 10^{-2}$ | $3.0 \times 10^{-2}$ | 1.02 |
| Transformer (“O → C”) | Shuffle (“O → C”) | $6.7 \times 10^{-4}$ | $1.4 \times 10^{-3}$ | 1.34 |
| Transformer (“O → C”) | Transformer (“C → C”) | $4.9 \times 10^{-1}$ | 2.9 | 0.07 |

**Table 1.** Statistics for ten comparisons in the main text. All comparisons were assessed using a one-sided Wilcoxon signed-rank test, with the null hypothesis that “Comparison 2 outperforms Comparison 1.” The $p$-values underwent Holm-Bonferroni correction for the six comparisons. The table reports both uncorrected and corrected $p$-values, along with effect sizes calculated using Cohen’s $d$. In the “Comparison 1 and 2” column, acronym abbreviations (O and C) represent Overt and Covert, respectively. For example, “O → O” signifies “Overt → Overt.”

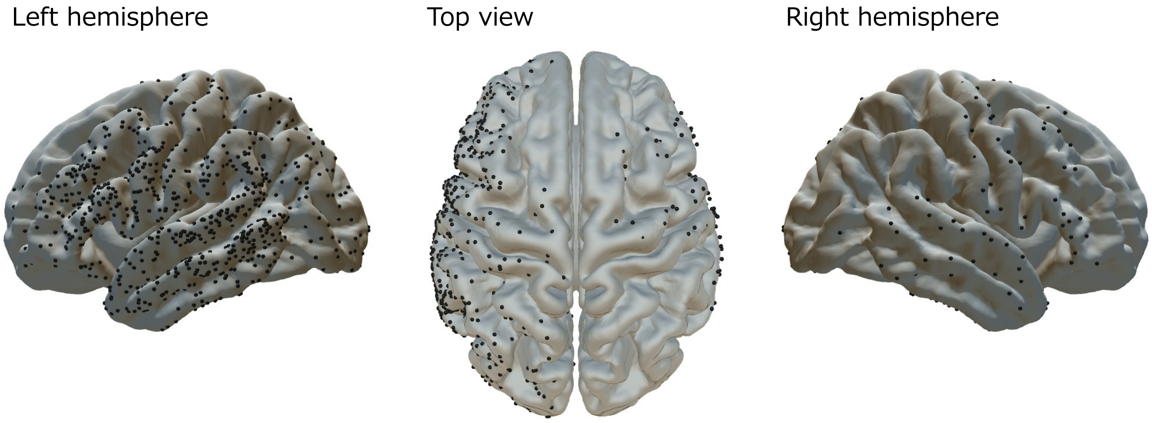


**Figure 1.** ECoG electrode coverage across all participants.

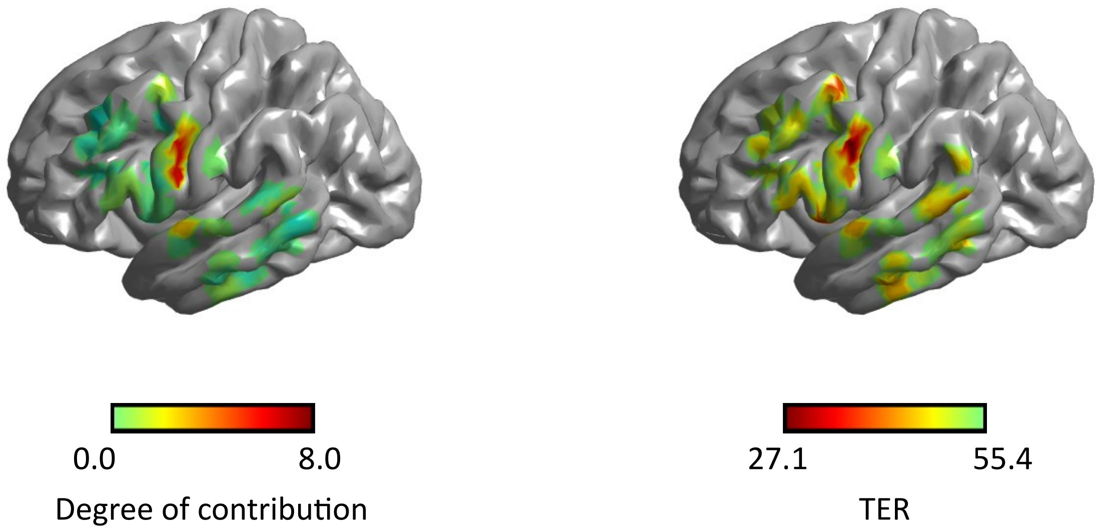


**Figure 2.** Comparison of the electrode contribution distribution and the TER distribution for js11 of "Overt → Overt." The TER distribution is obtained through training and evaluation using a single electrode. Each TER is computed based on 80 results from a 5-fold cross-validation, with 64 used for training and 16 for evaluation.

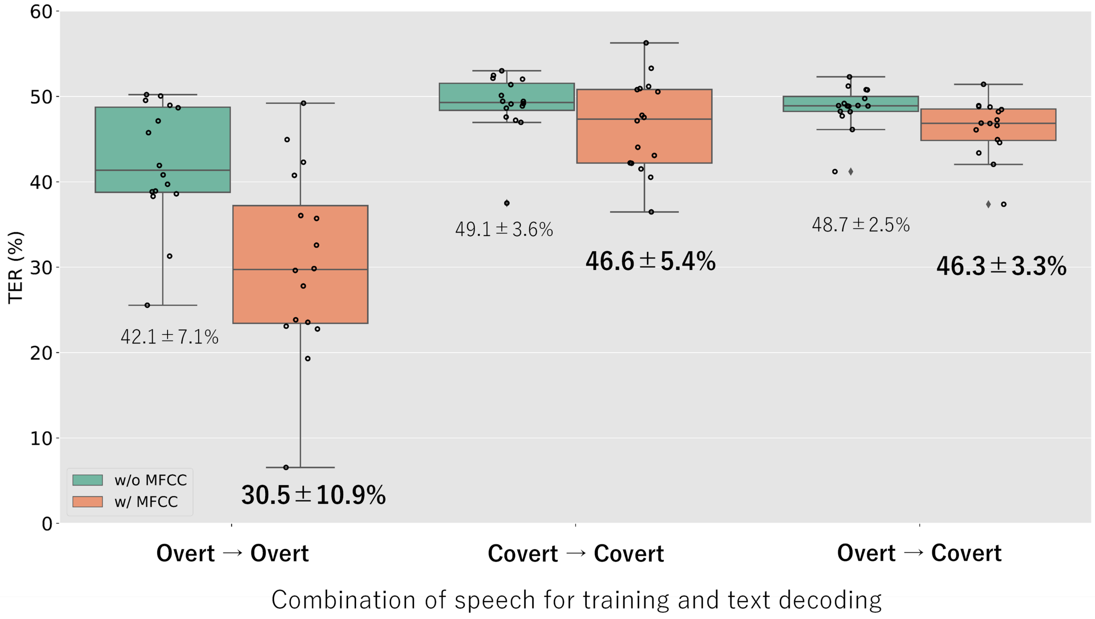


**Figure 3.** Decoding performance obtained by either using or not using MFCCs. The figure presents a boxplot depicting the distribution of TERs for three combinations of training and decoding. Each dot on the plot represents an individual participant. The green boxes represent the TER distribution of the Transformer with no MFCCs, whereas the orange ones represent the Transformer with MFCCs. We defined outliers as values that are 1.5 times the interquartile range (IQR) away from the median. The IQR represents the range of values within the middle 50% of the data. The numbers in the figure indicate the average ± standard deviation.

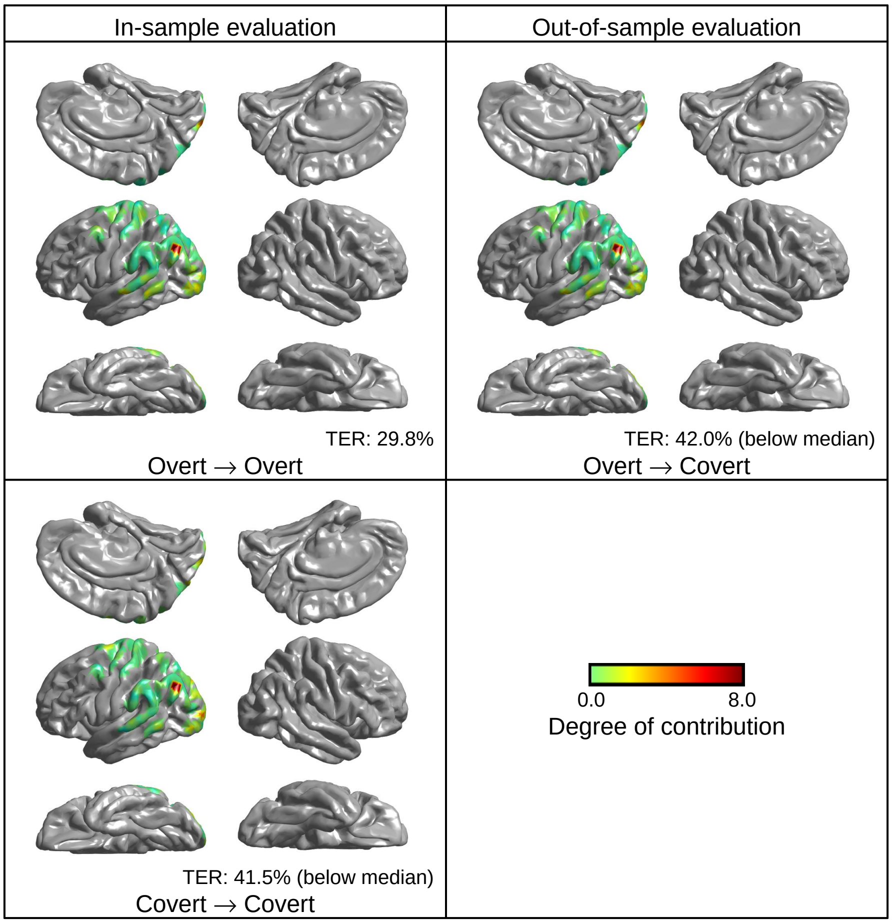


**Figure 4.** Electrode contributions for js1. TERs for js1 in "Covert → Covert" and "Overt → Covert" combinations are below the median among all participants. The inferior parietal cortex emerges as the most significant contributing area for all combinations.

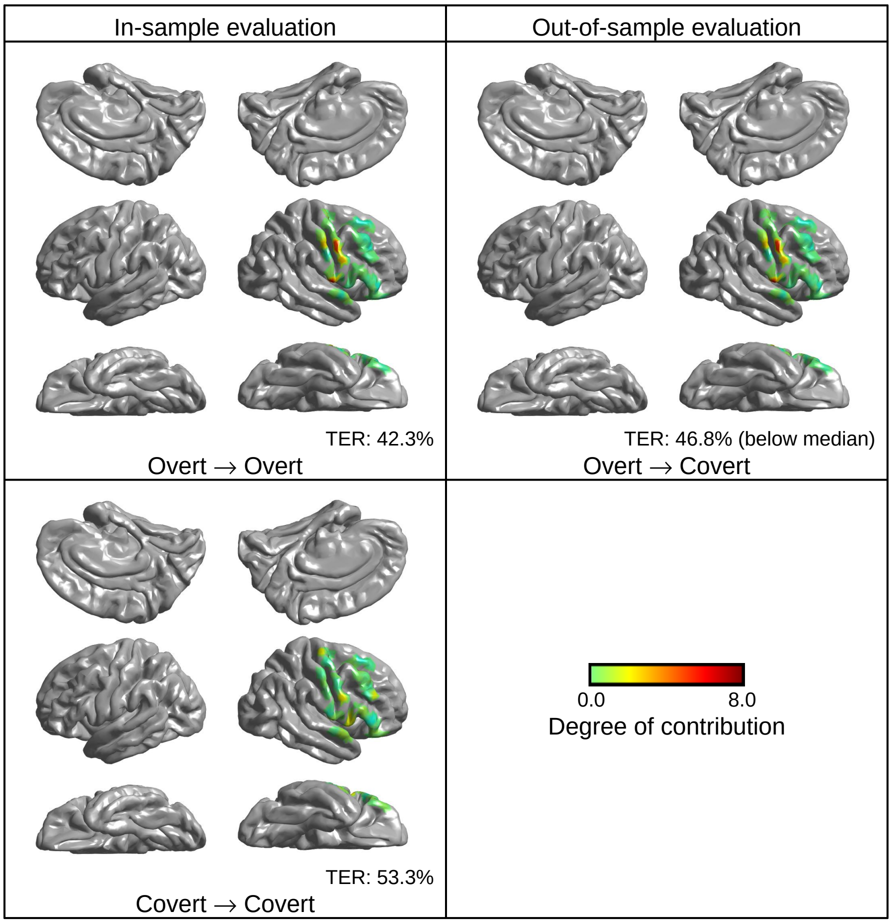


**Figure 5.** Electrode contributions for js2. Only the TER for js2 in the "Overt → Covert" combination is below the median among all participants. The SMC in the right hemisphere emerges as the most significant contributing area for this combination.

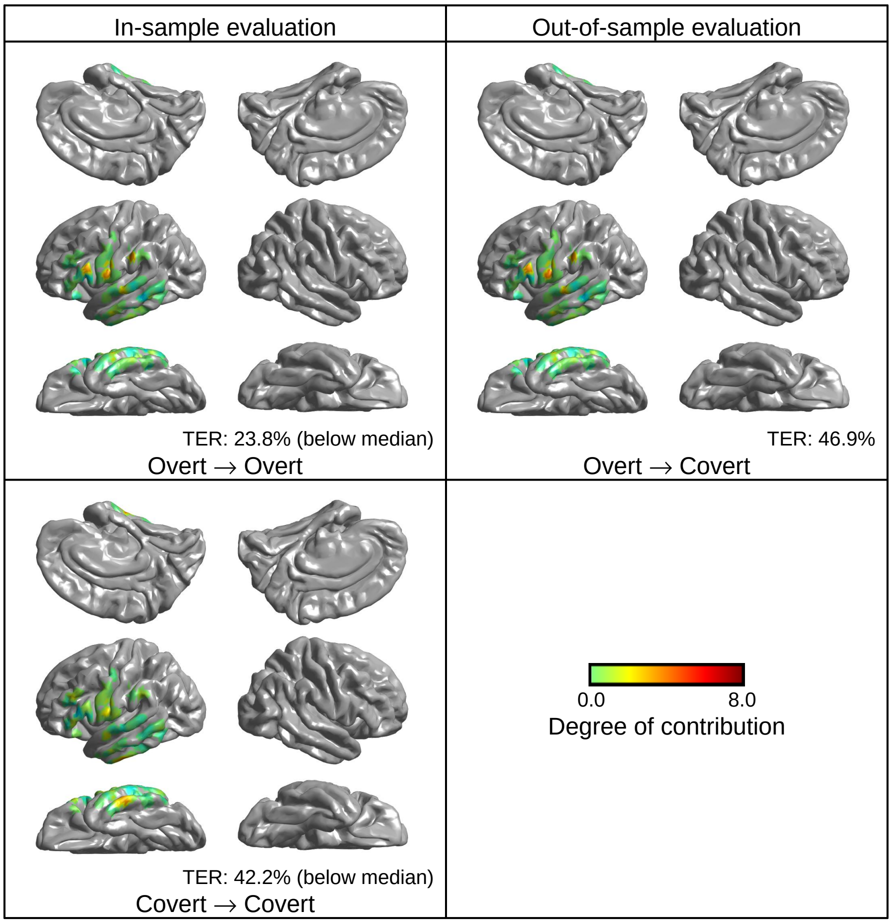


**Figure 6.** Electrode contributions for js3. TERs for js3 in "Overt → Overt" and "Covert → Covert" combinations are below the median among all participants. Broca's area, the SMC, and Wernicke's area for "Overt → Overt," emerge as significant contributing areas. The inferior temporal gyrus for "Covert → Covert" is a slightly contributing area.

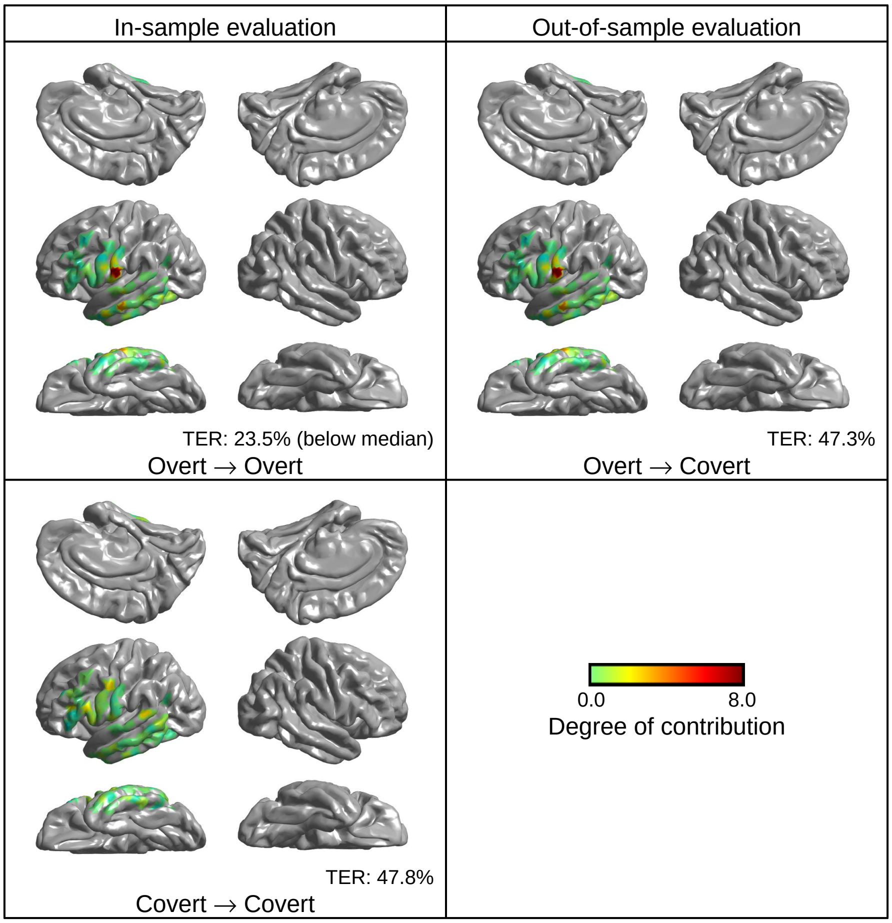


**Figure 7.** Electrode contributions for js4. Only the TER for js4 in the "Overt → Overt" combination is below the median among all participants. The SMC for "Overt → Overt" emerges as the most significant contributing area.

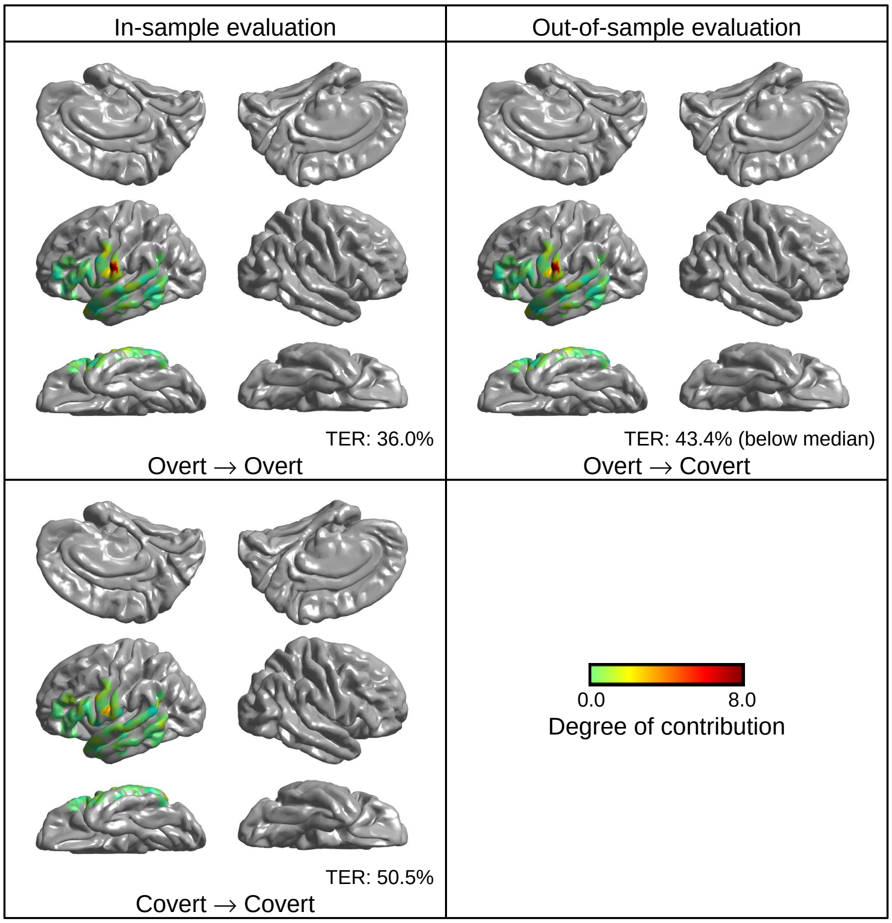


**Figure 8.** Electrode contributions for js5. Only the TER for js5 in the "Overt → Covert" combination is below the median among all participants. The SMC for "Overt → Covert" emerges as the most significant contributing area.

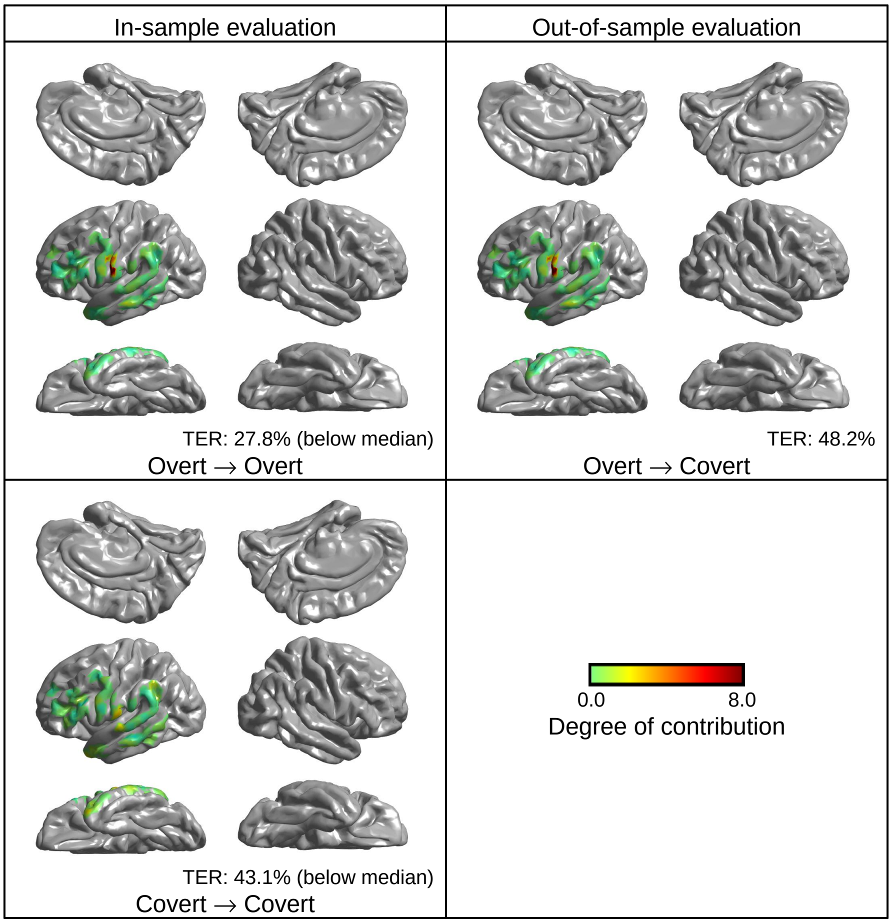


**Figure 9.** Electrode contributions for js6. TERs for js6 in "Overt → Overt" and "Covert → Covert" combinations are below the median among all participants. The SMC for "Overt → Overt" emerges as the most significant contributing area. No significant contribution area was found for "Covert → Covert."

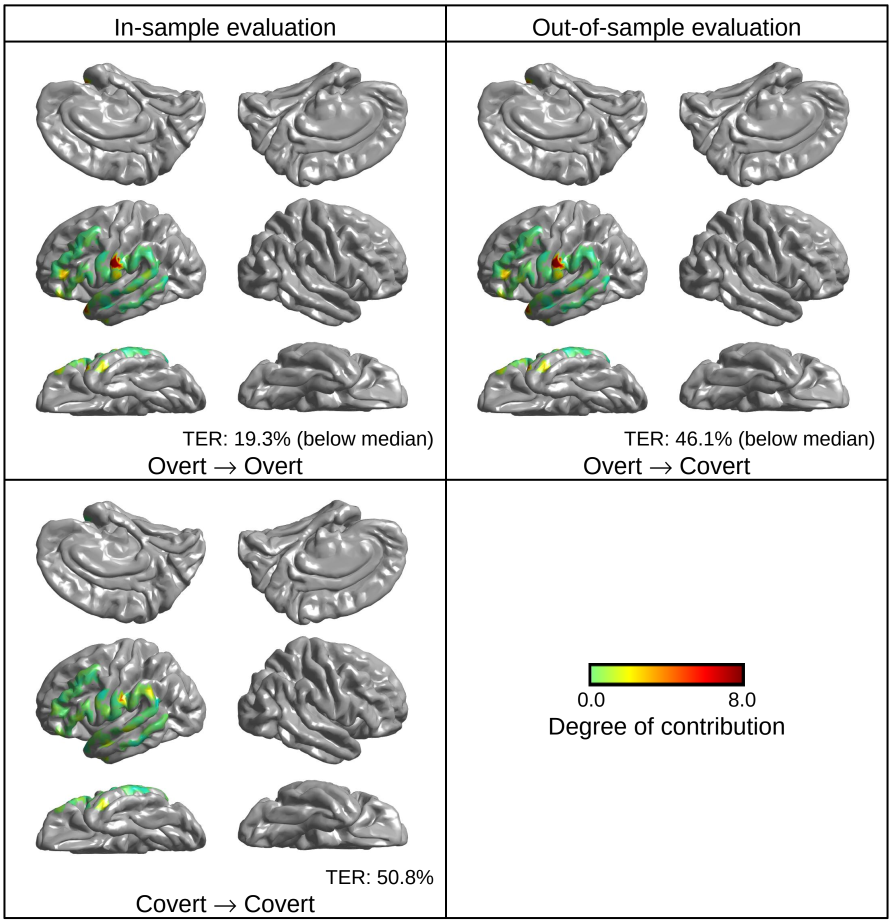


**Figure 10.** Electrode contributions for js7. TERs for js7 in "Overt → Overt" and "Overt → Covert" combinations are below the median among all participants. The SMC for these combinations emerges as the most significant contributing area.

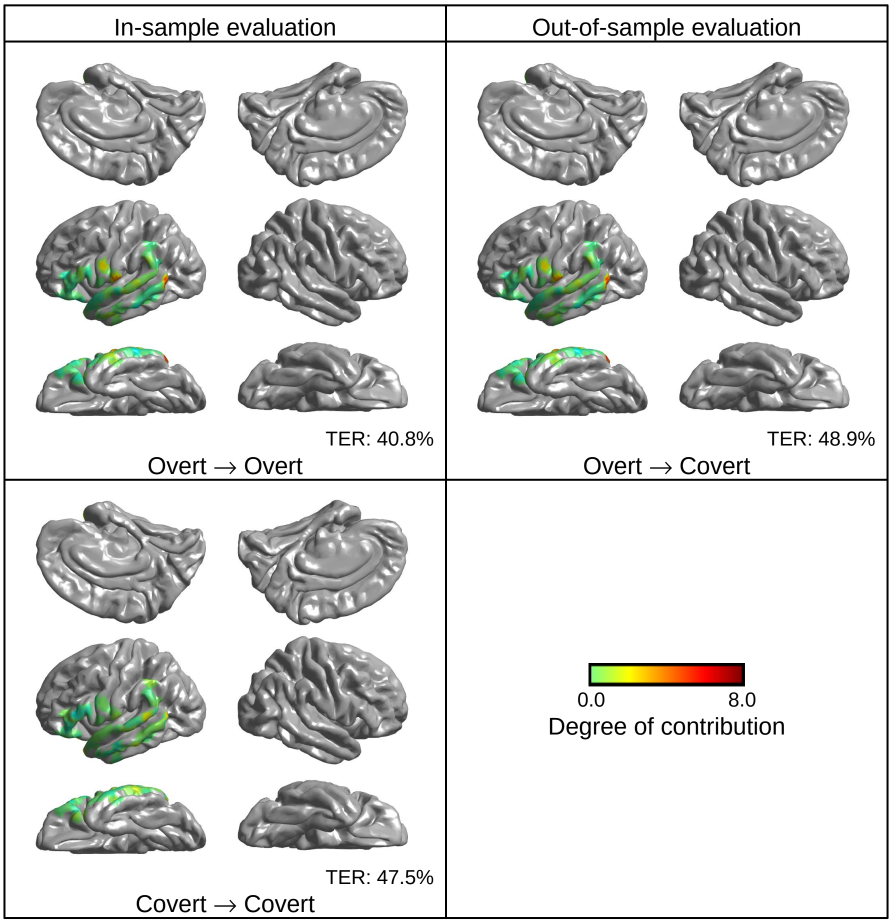


**Figure 11.** Electrode contributions for js8. TERs for js8 in all three comparisons are above the median among all participants.

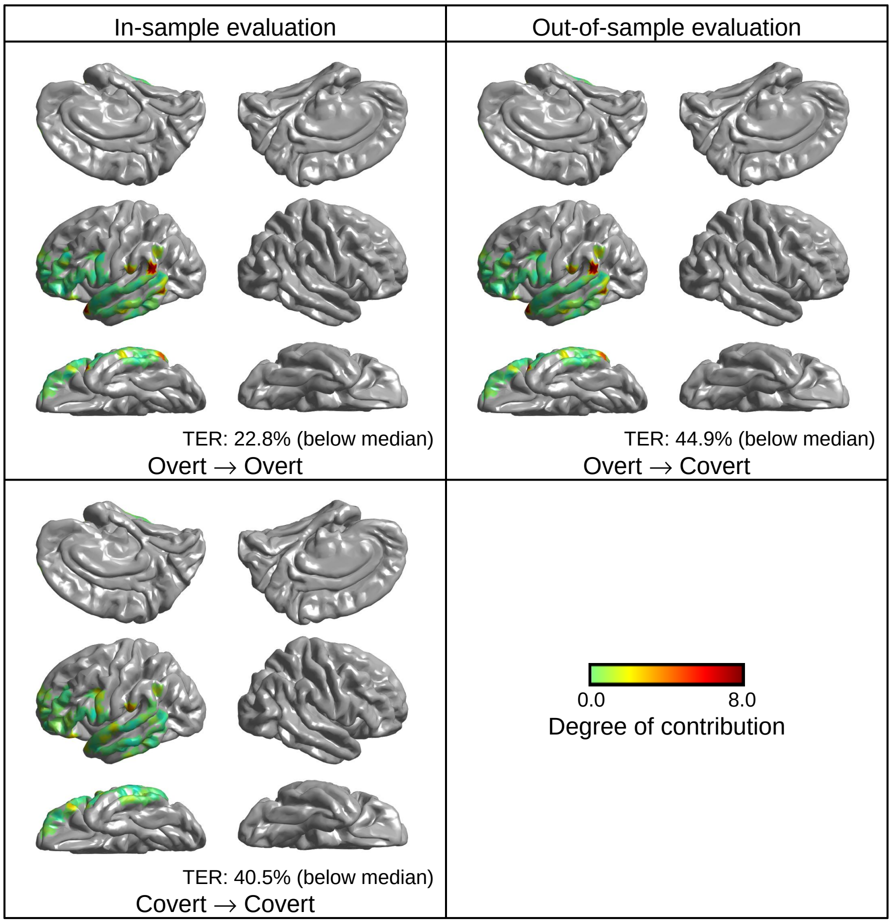


**Figure 12.** Electrode contributions for js9. TERs for js9 in all three combinations are below the median among all participants. Wernicke's area for both "Overt → Overt" and "Overt → Covert" combinations emerge as the most significant contributing area. No significant contribution area was found for "Covert → Covert."

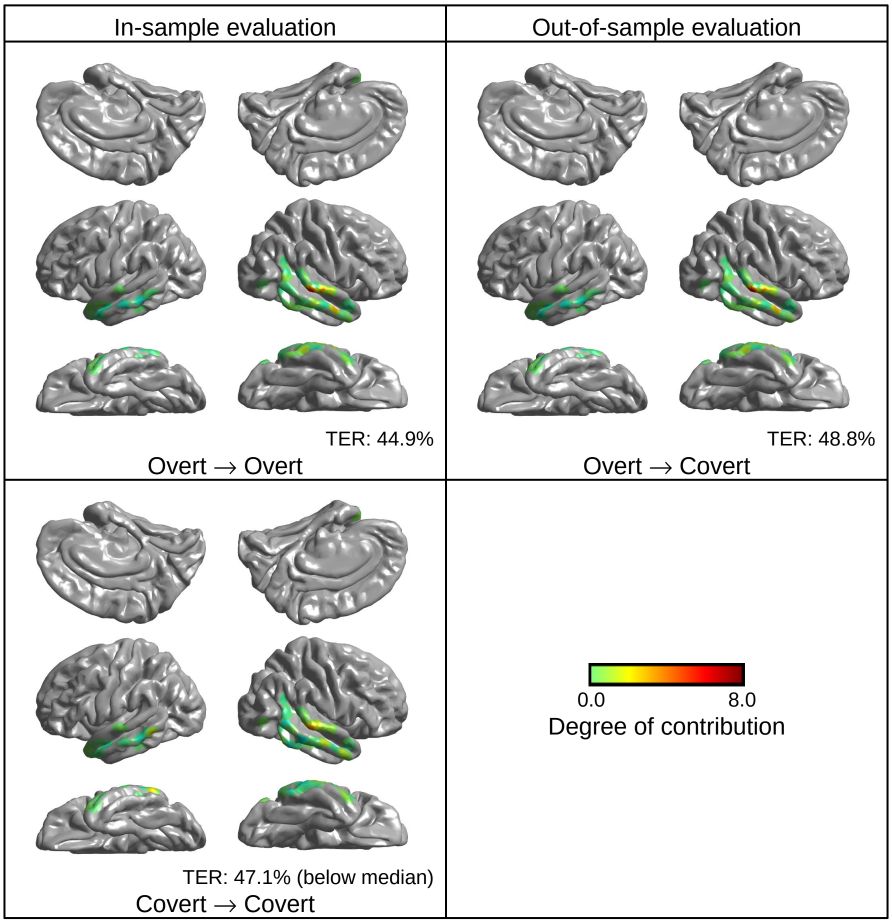


**Figure 13.** Electrode contributions for js10. TER for js10 in the "Covert → Covert" combination is below the median among all participants. The STG for the combination emerges as the slightly contributing area.

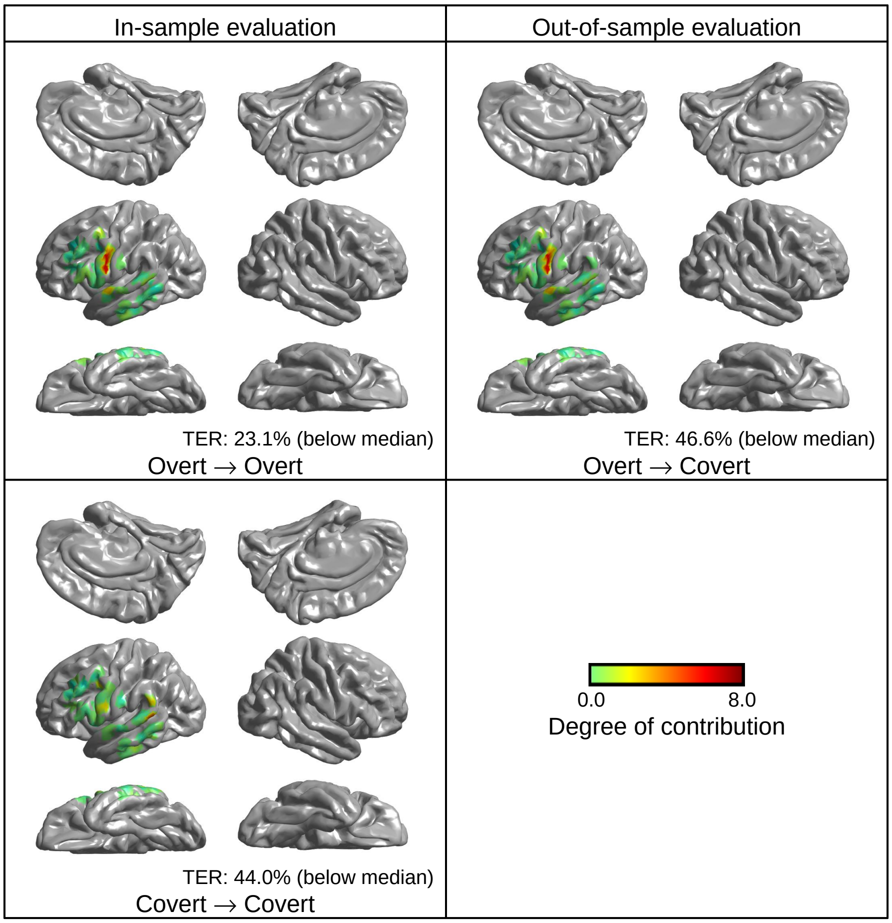


**Figure 14.** Electrode contributions for js11. TERs for js11 in all three combinations are below the median among all participants. The SMC for both "Overt → Overt" and "Overt → Covert" emerges as the most significant contributing area. No significant contribution area was found for "Covert → Covert."

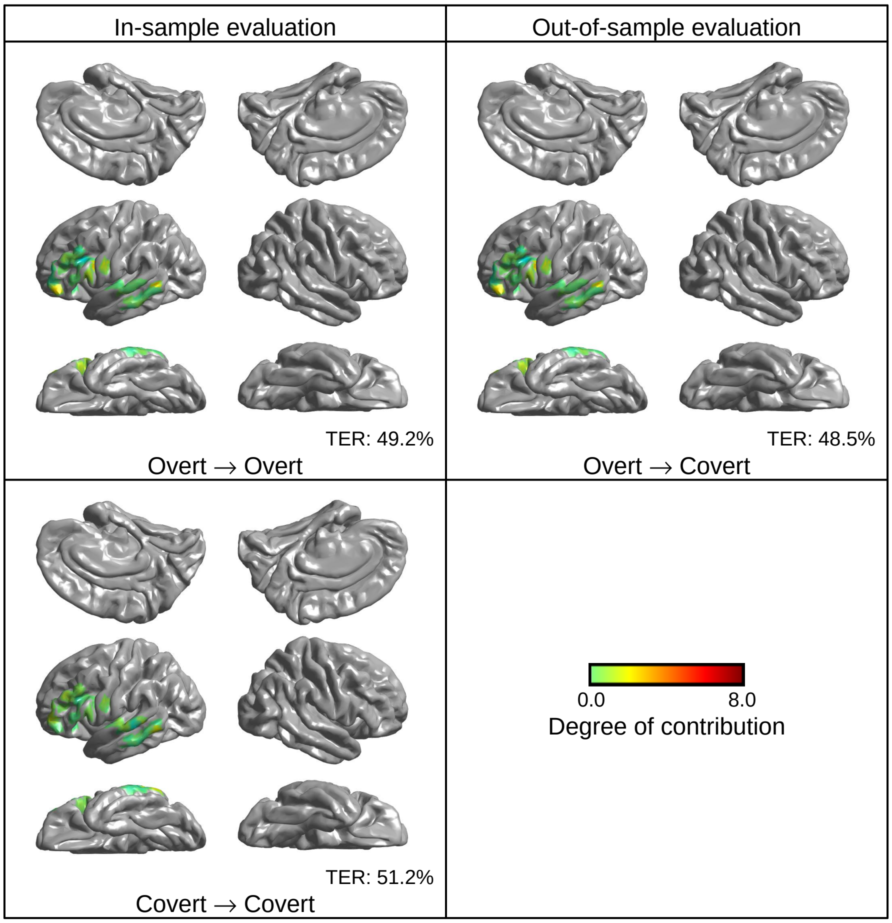


**Figure 15.** Electrode contributions for js12. TERs for js12 in all three comparisons are above the median among all participants.

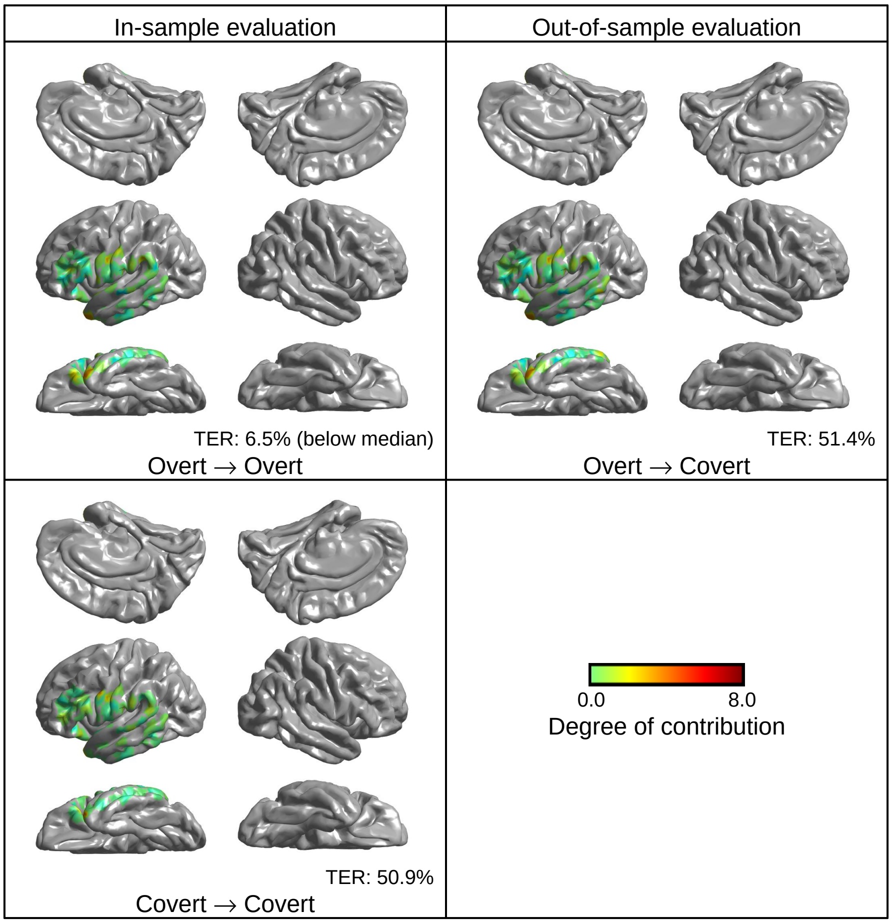


**Figure 16.** Electrode contributions for js13. TER for js13 in the "Overt → Overt," combination is below the median among all participants. The inferior temporal gyrus for "Overt → Overt" emerges as the most significant contributing area.

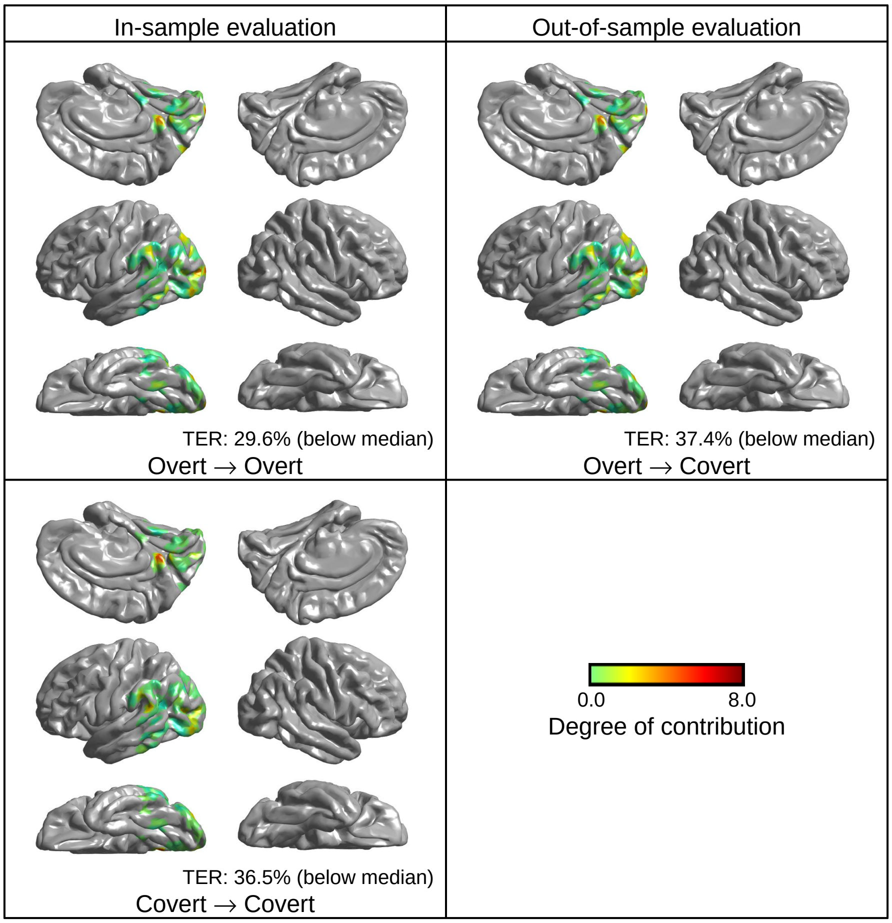


**Figure 17.** Electrode contributions for js14. TERs for js14 in all three combinations are below the median among all participants. The occipital lobe for these combinations emerges as the most significant contributing area.

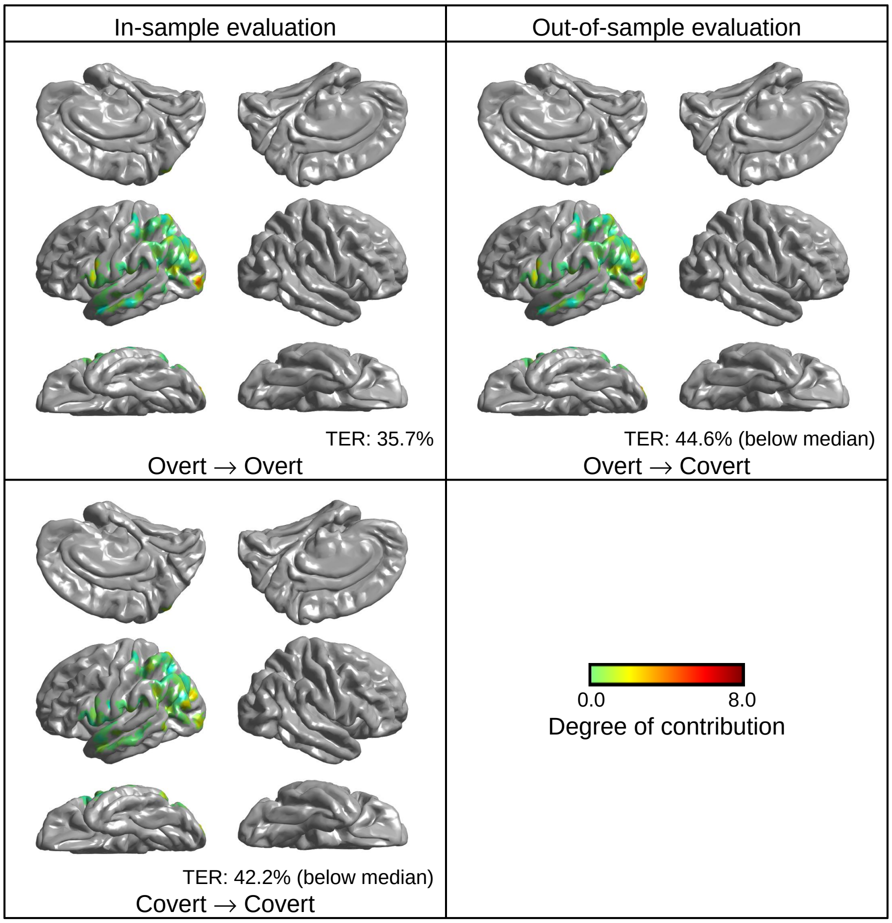


**Figure 18.** Electrode contributions for js15. TERs for js15 in the "Covert → Covert" and "Overt → Covert" combinations are below the median among all participants. The occipital lobe for "Overt → Covert" emerges as the most significant contributing area. No significant contribution area was found for "Covert → Covert."

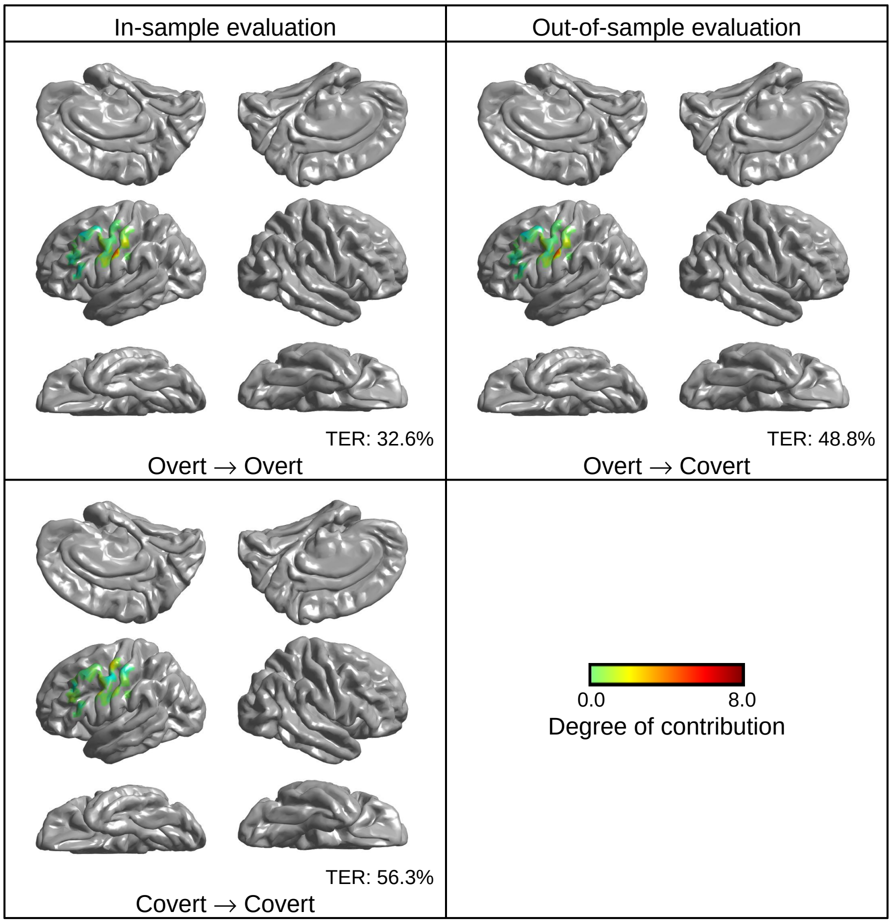


**Figure 19.** Electrode contributions for js16. TERs for js16 in all three comparisons are above the median among all participants.
